# Eye movement patterns under exposure to spatial disorientation illusions during simulated flight

**DOI:** 10.1101/2025.01.28.635199

**Authors:** Maya Harel, Idan Nakdimon, Oded Ben-Ari, Tom Schonberg

## Abstract

**Objective:** To identify eye movement patterns which are correlated with spatial disorientation (SD) events during flights in a flight simulator that induces SD.

**Background:** Spatial disorientation is one of the main causes of aviation mishaps. It results from illusions caused by misinterpreted vestibular or visual sensory cues, leading to an incorrect perception of an aircraft’s position, attitude, or motion. SD prevention is crucial, as there is no objective tool to detect its occurrence.

**Method:** Eye movements of 45 participants (30 aircrew members, 15 cadets) were recorded using Tobii Pro Glasses 2 in a Gyro-IPT SD flight simulator. Illusions were either vestibular or visual. Gaze metrics—such as fixations, saccades (rapid gaze transfers), and visits—were compared between participants who experienced SD and those who did not. Statistical analyses were conducted to identify significant differences.

**Results:** Among 284 flight profiles, 136 SD occurrences were recorded (48%). During visual illusions, participants who more frequently checked the instrument panel were more likely to avoid SD. In contrast, during vestibular illusions, participants who examined the head-up display (HUD) more frequently had a lower probability of experiencing SD.

**Conclusion:** Efficient SD mitigation requires task-specific eye-movement strategies: mitigating visual illusions requires increased focus on the instrument panel, while mitigating vestibular illusions involves greater engagement with the HUD, challenging current standard instructions.

**Application:** These findings may inform training programs aimed at improving performance in high-risk SD profiles. Additionally, the results support the development of real-time SD alert systems to help mitigate or prevent SD-related incidents.

**Precis:** This study examined eye-movement patterns during flight in a flight simulator to identify mechanisms underlying spatial disorientation (SD) and strategies for SD avoidance. Findings reveal that to avoid visual illusions greater instrument focus is required, whereas to avoid vestibular illusions, HUD engagement is beneficial. Results could be applied to training programs to enhance flight safety and reduce SD incidents.

## Background

Spatial orientation refers to an individual’s ability to correctly perceiving their position and motion relative to the earth. This perception is achieved through visual and vestibular cues or intellectually constructed from focal visual, verbal, or other symbolic data (Gillingham, 1992). In aviation, spatial orientation pertains to an aircrew member’s ability to recognize the aircraft’s velocity, altitude, heading direction, rotation, and pitch angle. **Spatial Disorientation (SD)** occurs when these elements are misperceived, leading to errors in interpreting motion, position, or attitude due to absent or misleading cues, primarily from the visual or vestibular systems (Cheung & Hofer, 2003).

SD in flight is a leading cause of aircraft mishaps attributable to operator error (Gillingham, 1992). It is responsible for 5 to 10% of all mishaps, 90% of which are fatal (Antunano, 2005). A U.S. Air Force report found SD contributed to 30% of mishaps from 1990 to 1999 and 26% from 1999 to 2009 (Gibb et al., 2011). Similarly, SD was a factor in 20% of fatal mishaps in the U.S. Navy between 1990 and 2000 and 38% between 2000 and 2017 (Meeks et al., 2024). SD-related mishaps are frequently caused by undetected illusions, with 80% of incidents occurring without aircrew awareness (Gregg, 1955)

Several visual and vestibular illusions that can occur during flight pose a risk for SD in aircrew members. The main illusions are described as follows:

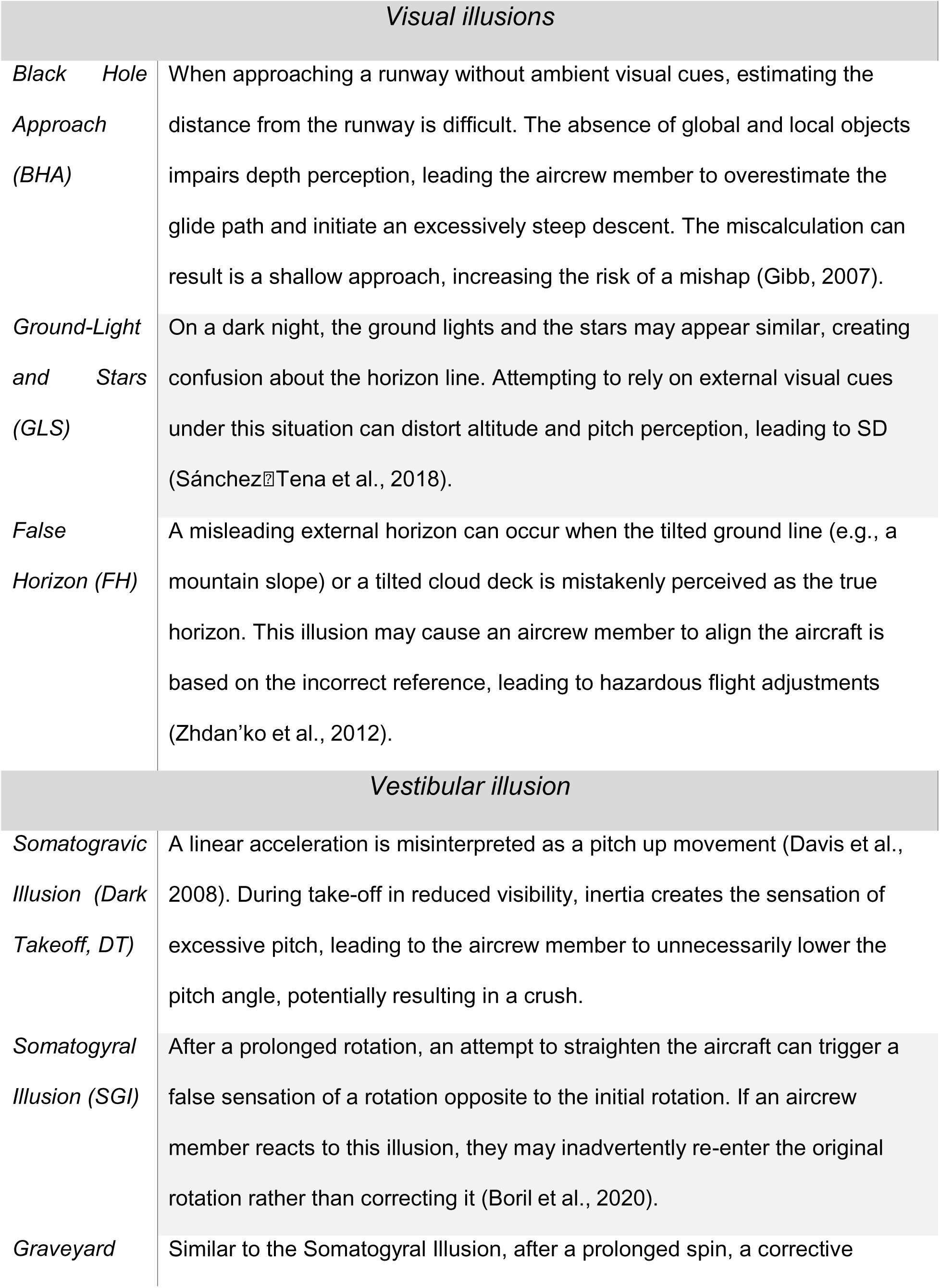
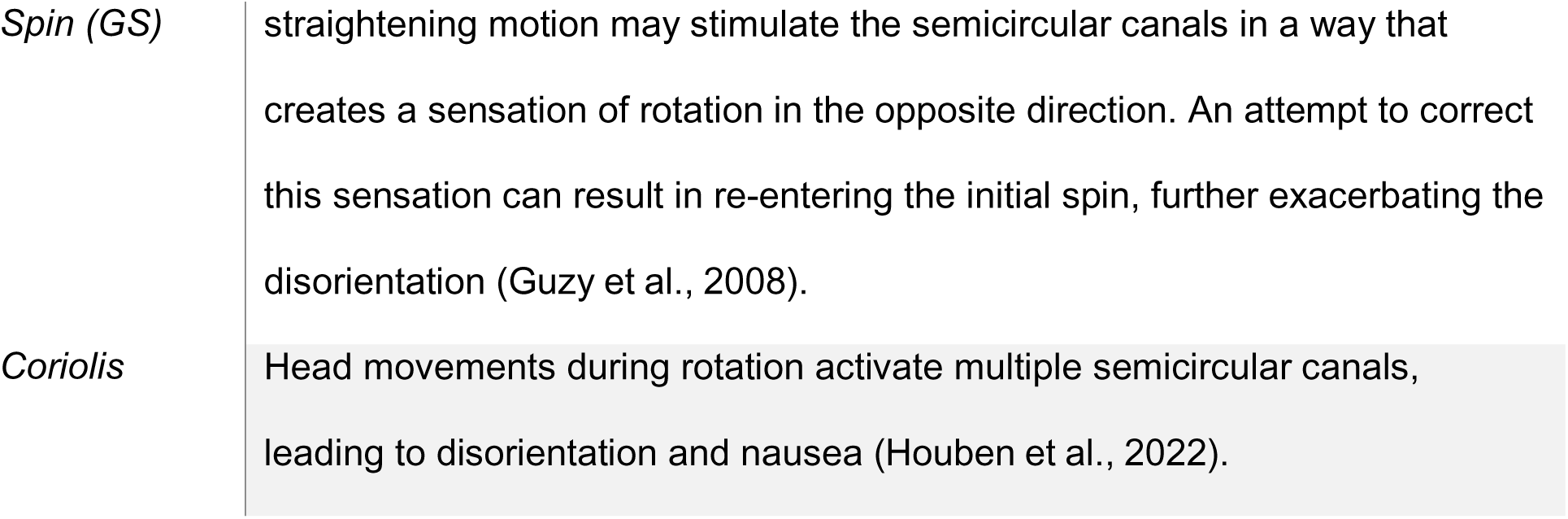

High-fidelity simulators replicating SD-inducing conditions are recommended for SD training (Cheung & Wong, 1998). An SD simulator is a flight simulator specifically designed to induce and study spatial disorientation by integrating both visual and vestibular stimuli: (1) Visual stimuli – The simulator includes screens that replicate the aircraft’s external environment, simulating real-world flight conditions. Additionally, instrument panels are displayed, providing an objective representation of the aircraft’s state for the aircrew member. (2) Vestibular stimuli – The simulator incorporates motion capabilities across the three principal axes of movement: pitch, roll, and yaw. The aircrew member actively controls these movements using the throttle lever and flight control stick, creating a dynamic training experience. An SD simulator integrates visual and vestibular stimuli to replicate flight conditions, enhancing spatial disorientation training. These simulators allow researchers to examine physiological and perceptual changes that occur during SD events.

Studies have demonstrated that vestibular stimulation can influence heart rate and respiration (Ilbasmis & Yildiz, 2017). Research has also shown that nighttime climbing perception is affected by optokinetic and cervico-ocular reflexes (Tamura et al., 2016; Maurice & Gioanni, 2004). Additionally, participants consistently perceive roll tilt during a turn, yet their pitch perception during takeoff is slower and less consistent (Tribukait et al., 2013). Furthermore, when no visual cues were available, attempts to straighten after a prolonged turn often led to an involuntary return to the original turn direction (Nooij & Groen, 2011).

Eye-tracking technology provides valuable insights into pilots’ visual scanning patterns, revealing their attention to critical instruments and external cues. *Fixations*, indicate sustained attention (Valliappan et al., 2020; Unsworth et al., 2020) and may reflect thorough cognitive processing of critical information (Just & Carpenter, 1976), or, conversely, suggest cognitive overload (Hauland, 2019). Research has shown that prolonged fixations on critical instruments were associated with better situational awareness, as pilots dedicated more time to interpreting vital information needed for safe flight operations (Goldberg & Kotval, 1999). Meanwhile, the *Visit* measure provides a broader perspective on how often a pilot re-engages with complex visual stimuli, suggesting the re-visited areas contain necessary information for decision making (McLaughlin et al. 2017).

*Saccades*, or rapid eye movements between fixations, are crucial for active scanning (Barfield & Furness, 1995). Frequent and large saccades indicate active situational awareness, whereas reduced saccadic activity may signal a narrowed focus or cognitive tunneling, where a pilot becomes overly reliant on a single instrument and may neglect other critical information (Di Stasi et al., 2013). Additionally, mental effort (Joshi et al., 2016), and unexpected sensory outcomes (Yokoi & Weiler, 2021).

Thus, eye-tracking can help us understand how aircrew members respond to situational changes, such as SD illusions. Eye-tracking serves as both an objective physiological response indicator—providing information about saccades, and visual movement patterns—as well as a subjective indicator of experience, as fixation areas reveal which areas of interest received attention. When examining visual movement patterns in pilots using imagery that simulated the external visual environment of the cockpit, research found that efficient movement across multiple areas of interest (AOIs), supported by well-timed saccades, was crucial for maintaining situational awareness and preventing SD (Thomas & Wickens, 2016). Similarly, prolonged fixations and visits on critical instruments were associated with better situational awareness, as pilots spent more time interpreting vital information needed to maintain safe flight operations (Goldberg & Kotval, 1999).

Research conducted in the Korean Air Force examined differences between movement patterns across flight instruments between a group where a verbal report of their flight data was required and one that wasn’t (Kang et al., 2021). That study found that the verbal report group focused more on specific areas of interest (AOI) and had a higher fixation frequency in the altitude measure in the Head-Up Display (HUD) in both types of SD: Vestibular (Coriolis, GS) and Visual (GS). A Japanese Air Force study examined eye movements during DT illusions found an automatic downward motion during the pitch-up motion, a reflexive response meant to stabilize the visual input (Tamura et al., 2016). When comparing experienced pilots and non-pilot participants in Vestibular illusions (FH, SGI, and Coriolis), research found that experienced pilots were as susceptible to SD as non-pilots (Bałaj et al., 2018).

Previous studies focused on a limited range of AOIs and illusions although a broader range of scenarios is required to gain a more comprehensive understanding of SD mechanisms. As Cheung (2013) claims, there is a need for renewed training that will improve pilots’ physical and mental performance during sensory conflict (Bob Cheung, 2013). Previous studies have also focused only on one aircrew member population. Comparing experienced pilots to navigators and cadets can help us create better learning paradigms to help prevent SD events. A broader model including different populations can add a data-driven angle to identify underlying mechanisms.

Therefore, based on existing literature, we aimed to use eye-tracking to study the mechanisms leading to SD events. Specifically, we aimed to compare eye-movement patterns of aircrew members during both visual and vestibular illusions that have the potential to create an SD event. To simplify the multiple variants of analysis we focused on several factors: role (pilot and navigator), experience (cadets vs. experiences pilot) and two main AOIs (HUD vs. instruments). This aimed to separate aircrew members who experienced the SD effect from those who maintained correct situational orientation. The overreaching goal was to lay the foundations to create more targeted training to mitigate causes of SD events, using the objective tool of eye-tracking.

## Methods

### Study Design

We collected participants as part of their pre-existing training program. The participants were from two groups: flying cadets and experienced pilots. The study was approved by the Ethics Committee of the Israeli Defense Forces based on the ICH-GCP principles. This is an exploratory study and was not pre-registered, yet the procedure and sample size were predetermined by the ethics protocol.

### Apparatus and measurement

#### Eye-tracking glasses

The Tobii Pro Glasses 2 Live View Wireless 100 were used to collect data, the parameters extracted from the glasses are elaborated in *Table 1* in the variables section.

**Table 1.**
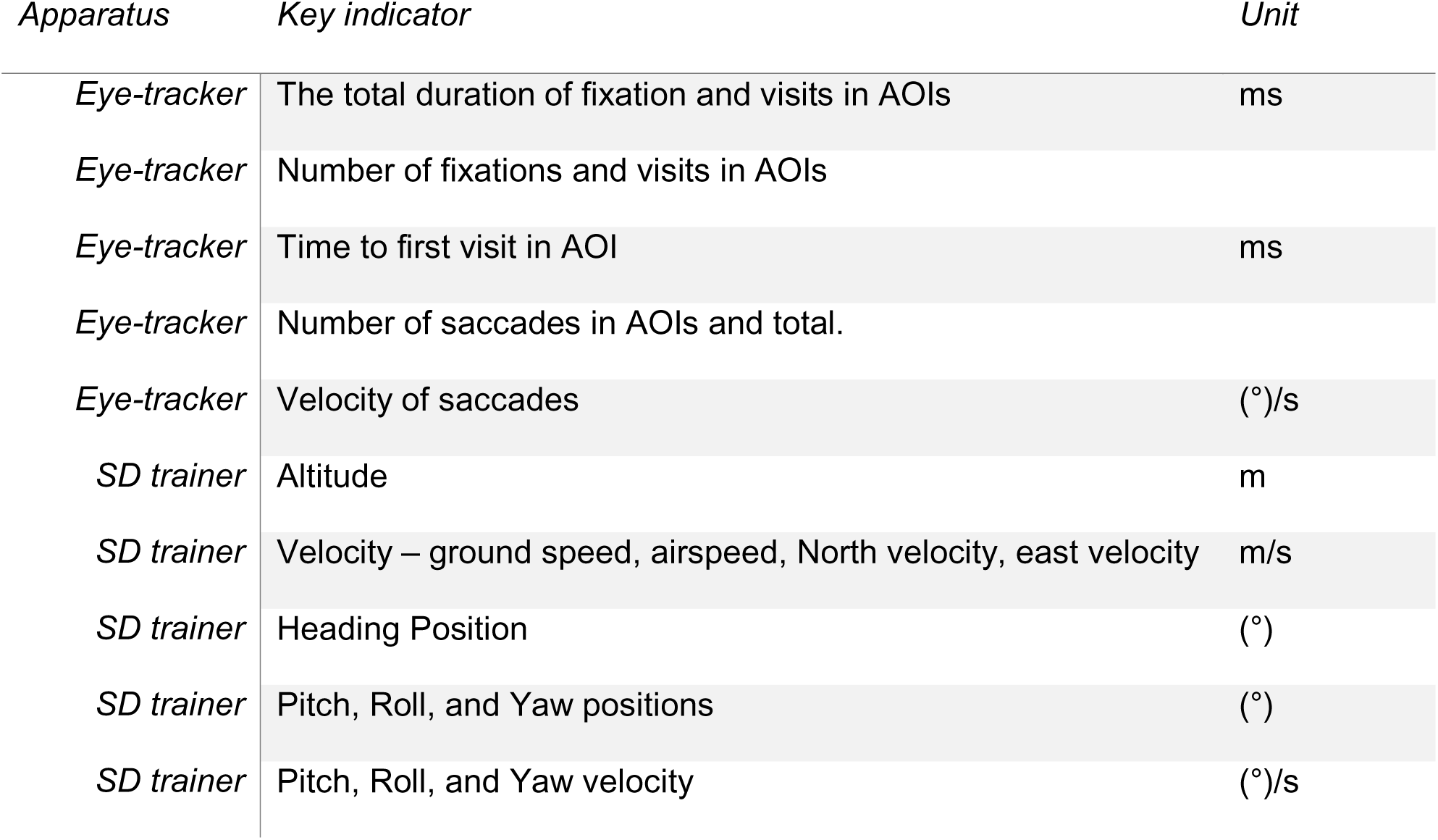
Eye-tracking and SD trainer parameters.

| *Apparatus* | *Key indicator* | *Unit* |
| --- | --- | --- |
| *Eye-tracker* | The total duration of fixation and visits in AOIs | ms |
| *Eye-tracker* | Number of fixations and visits in AOIs |  |
| *Eye-tracker* | Time to first visit in AOI | ms |
| *Eye-tracker* | Number of saccades in AOIs and total. |  |
| *Eye-tracker* | Velocity of saccades | (°)/s |
| *SD trainer* | Altitude | m |
| *SD trainer* | Velocity – ground speed, airspeed, North velocity, east velocity | m/s |
| *SD trainer* | Heading Position | (°) |
| *SD trainer* | Pitch, Roll, and Yaw positions | (°) |
| *SD trainer* | Pitch, Roll, and Yaw velocity | (°)/s |

#### SD trainer

We used The Gyro-IPT simulator (Environmental Tectonics Corporation, Inc., Southampton, US), located at the Israeli Air Force Aeromedical Center. This dynamic motion-based simulator has 3 degrees of freedom (roll ±25°, pitch ±25°, yaw 360°, and Heave =/− 25cm) and has an out-the-window visual display (120cm * 40cm 3 windows wide field of view). The flight instruments panel displayed in the cabin represents typical indicators that are applied in the pilots’ aircraft (Military Institute of Aviation Medicine, Warsaw, Poland et al., 2023).

### Participants

Aircrew members (n = 30): operational experienced high-performance aircrew members: pilots and navigators, 30-50 years old (average 37.74±5.46 years, 1 SD), with 12-22 years of experience.

Flying cadets (n = 15): pilot cadets and navigator cadets, in their second year of the flying academy (average age 19.84±0.01 years, 1 SD). The training profile for cadets included only 4 out of the 7 illusions, 2 visual (GLS, BHA) and 2 vestibular (SGI, and Coriolis) in the aircrew member training profile, as elaborated **Error! Reference source not found.**.

### Variables

The demographic parameters: Aircraft type (F - 15, F - 16, or T - 6), Aircrew member role (navigator or pilot), and age were collected and are reported below.

Eye-tracking parameters were extracted from the eye-tracking glasses using the Tobii Pro lab program. Flight parameters were extracted from the Gyro IPT program per flight profile. The data consisted of the aeromodel parameters (*Table 1*).

### Procedure

Forty-five participants were recruited, thirty active aircrew members and fifteen flying cadets. They were approached when they arrived at the SD simulator according to the Israeli Air Force training regulations. Prior to the beginning of the training session, they were presented with the study’s aims and the procedure. They were explained the study would not change the course of the mandatory training they arrived for and would only add the eye-tracking glasses. Those who agreed to participate signed the informed consent form. Then, a briefing session was provided for each participant about the simulator and the training procedure. The participants were equipped with the Tobii glasses, and a calibration process was conducted using the internal calibration algorithm of the Tobii Glasses 2. During the calibration process, the participants had to wear the Eye-tracking glasses while focusing on the center of the calibration target (*Figure* 2 - calibration target) located about one meter from the participant

**Figure 1.**
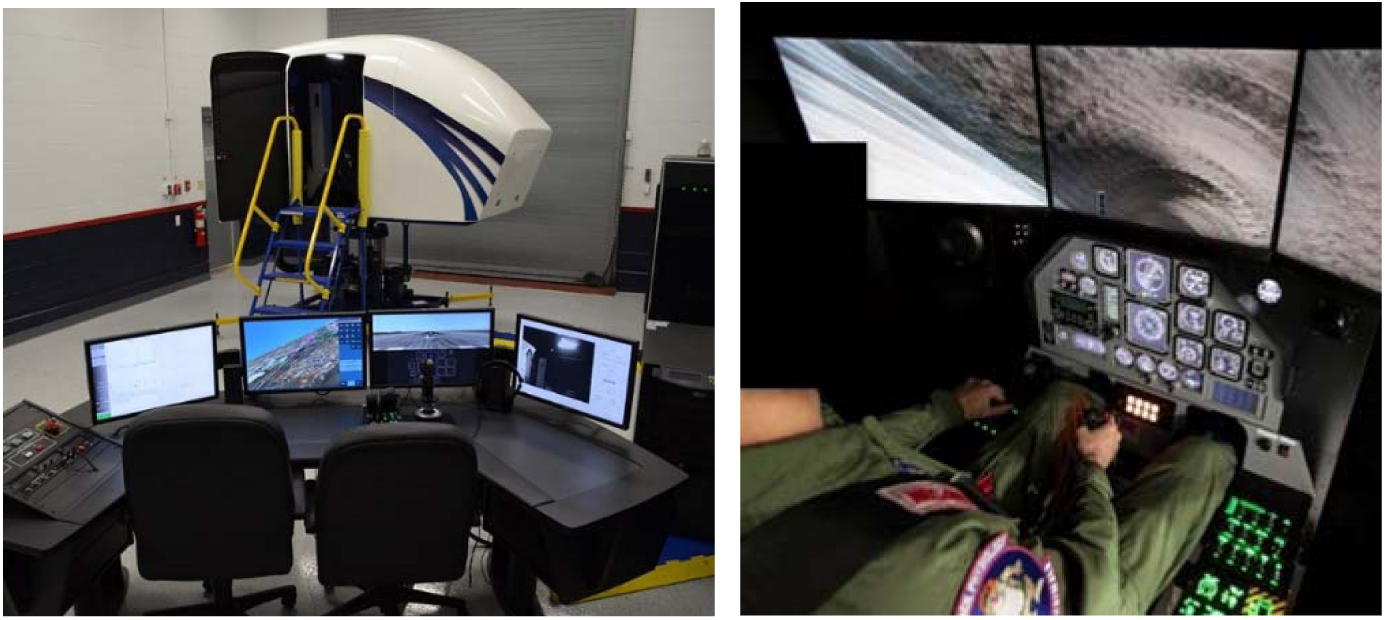
SD simulator. (1) Left: the SD simulator from the outside, which has a motion ability in 3 axes. (2) Right: internal view of the SD simulator containing a visual external display and the cockpit instruments panel.

**Figure 2.**
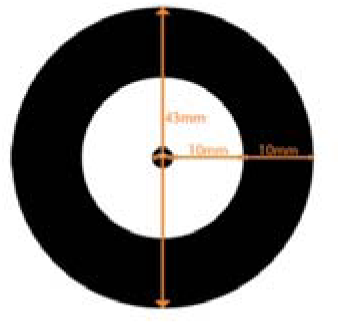
calibration target.

Then the participants entered the SD trainer and underwent the training procedure, which included up to 8 flight profiles, elaborated in the supplementary material. All subjects underwent the same illusions ordered as presented in the table. Each profile lasted 1 to 3 minutes, between profiles the subjects remained inside the SD trainer while the instructor debriefed about the SD illusion they experienced and for subjects who showed SD response explained what they should have done differently. The training procedure is summarized in *Figure 3*.

**Figure 3.**
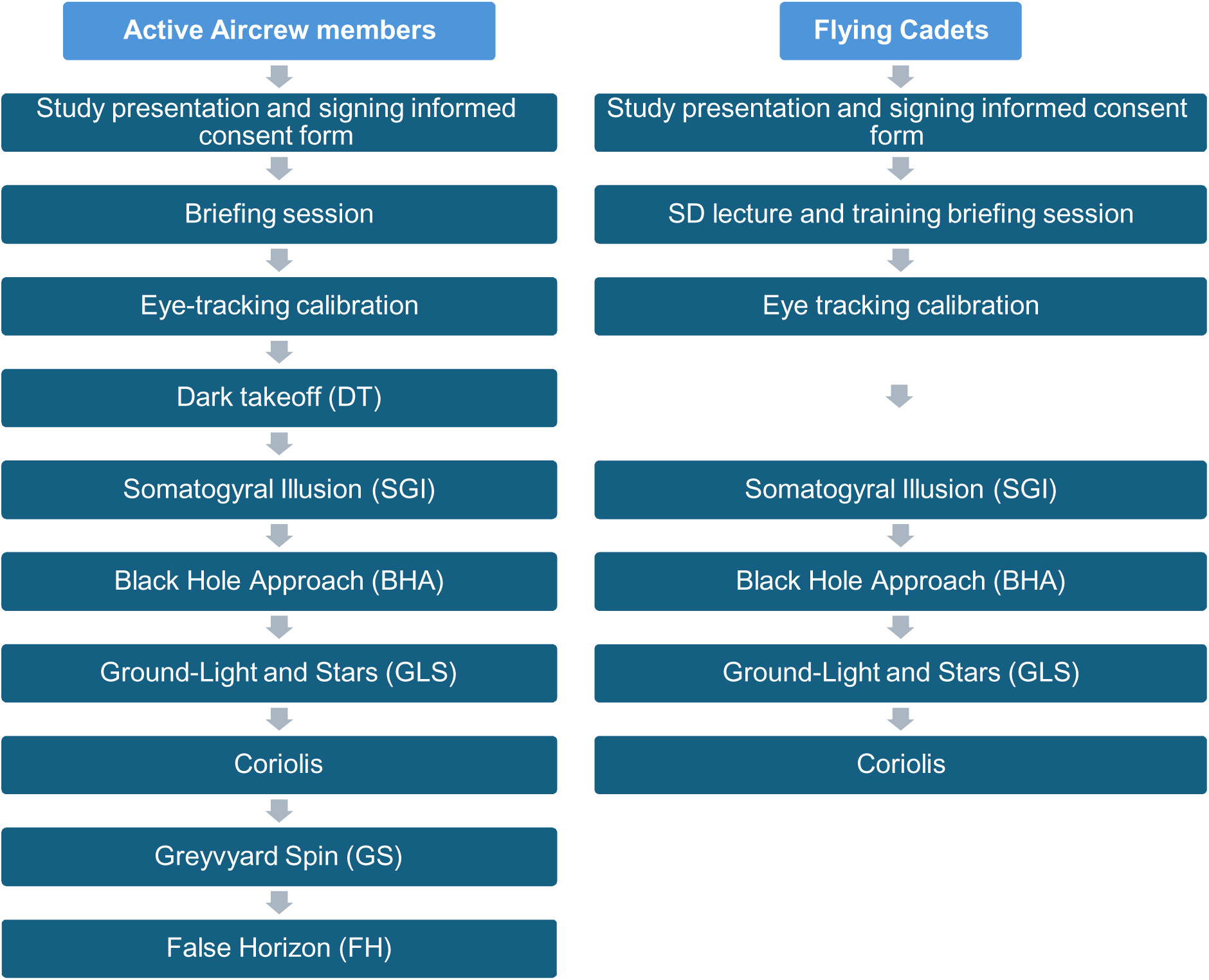
training procedure.

For each illusion the training profile, the SD window, criteria for identifying SD response, and a visual example of the profile timeline are presented in the supplementary material.

### AOIs

An AOI was marked for each instrument, HUD element, and external aircraft display (*Figure 4*):

**Figure 4.**
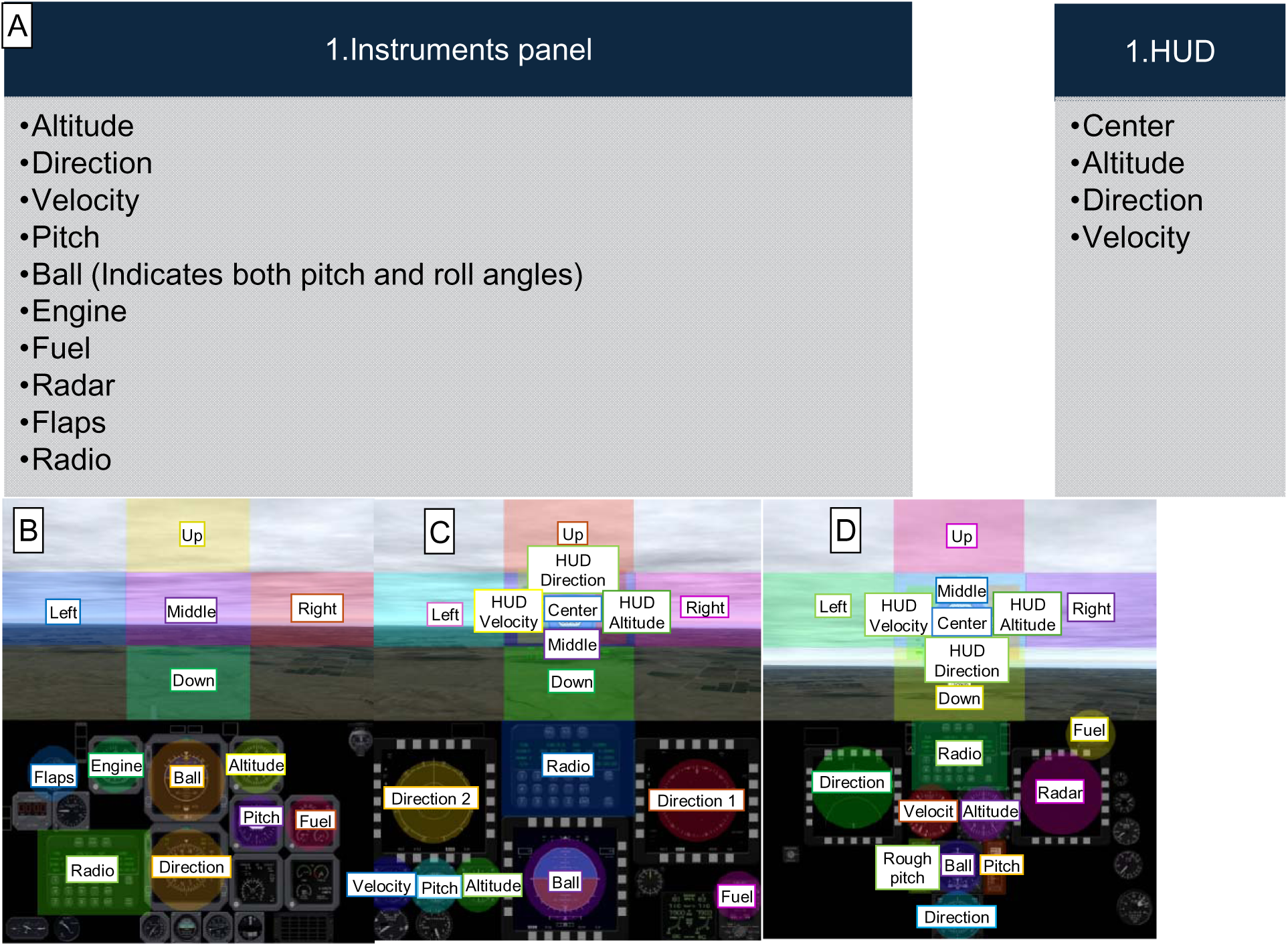
AOI list and mark over aircraft’s display. (A) The list of AOIs divided by location (B) The T-6 aircraft display of instruments, it does not have an HUD display (C) The F-15 aircraft display of the instruments panel, and the HUD. (D) The F-16 aircraft display of the instruments panel, and the HUD.

### Statistical analyses

The initial analysis of the eye-tracking data was performed using Tobii Pro Lab (Tobii PRO LAB), which produced the eye-tracking parameters outlined in *Table 1* for each Time of Interest (TOI) and Area of Interest (AOI) (*Figure 5*). These parameters were then entered into Python 3.11, where logistic regressions predicting SD response were conducted. Given the multiple comparisons between SD and non-SD groups across various illusions and AOIs, statistical significance was set at 0.05, and the Holm-Bonferroni multiple comparison correction was applied.

**Figure 5.**
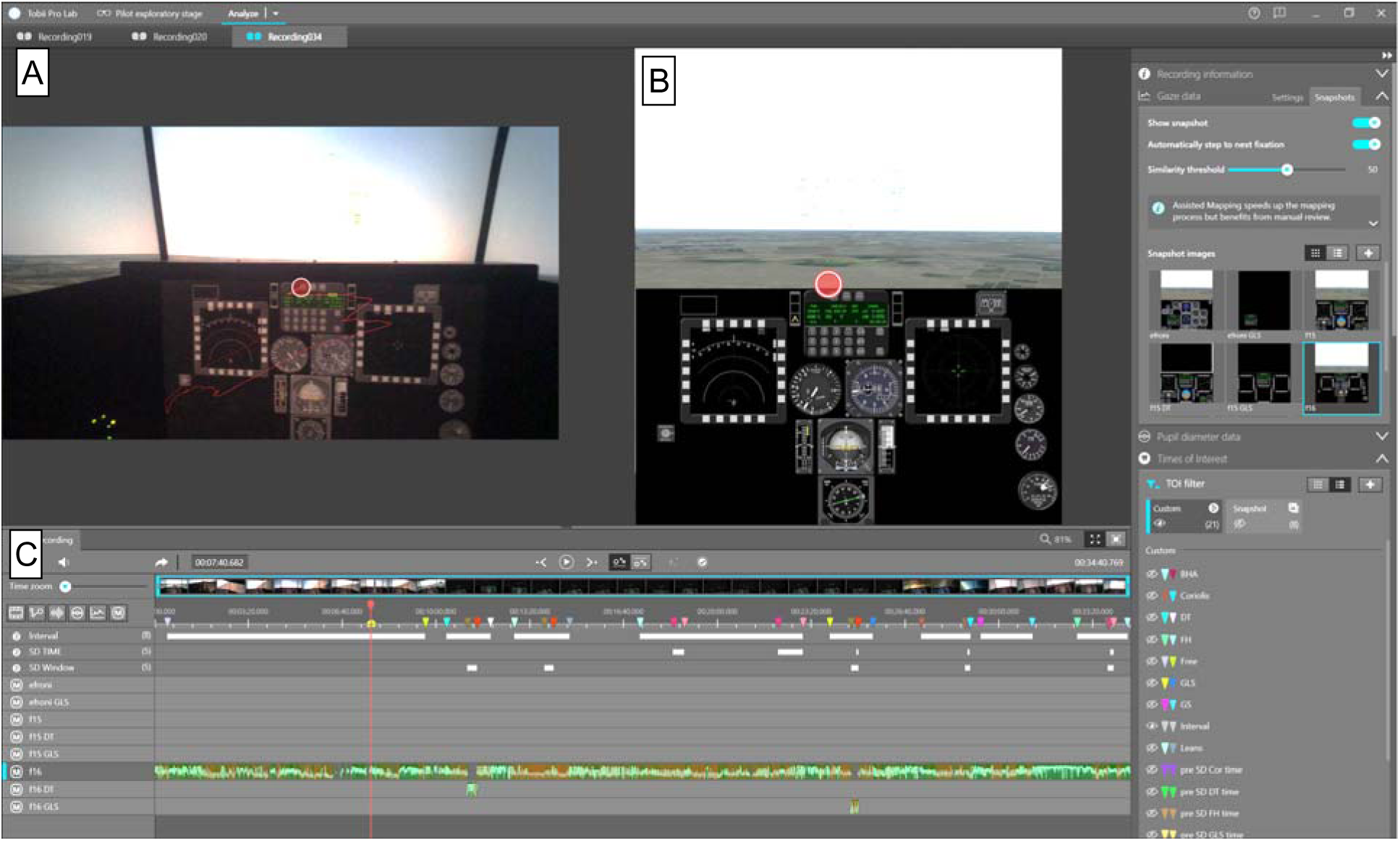
Tobii PRO Lab. The Tobii PRO lab statistical analysis program: (A): Video recording from the eye-tracker. (B): A reference snapshot for the compatible aircraft display. (C): Recording timeline with the following marks (up to down) - relevant event marks, interval, SD time, and TOI marks, gaze mapping from video to snapshot affinity.

The data analysis code and the row data are attached in the github repository: https://github.com/mayahar/Harel-et-al

## Results

### Demographics

Data from active aircrew members and flying cadets were divided into pilots and navigators. (*Table 2*).

**Table 2.** participants demographics. Position Aircraft n Percentages out of entire sample Average age Number of flight profiles Within the navigator cadets: all 5 participants performed the 4 profiles (summing to 20). Within the pilot cadets all 10 participants performed the 4 profiles and two performed BHA twice, (summing to 42 sorties). Within the F-15 navigators two of the participants performed all 7 profiles, one performed BHA twice, one performed DT twice, and one performed only 4 profiles (DT, SGI, GLS, and BHA) due to an eye-tracker malfunction (summing to 20 sorties). Within F-15 pilots all four participants performed all 7 profiles (summing to 28 sorties). Within F-16 navigators all ten participants performed the 7 profiles, seven performed BHA twice, and four performed DT twice (summing to 81 sorties). Within F-16 pilots All thirteen participants performed the 7 profiles, and one performed DT twice (summing to 92).

| *Position* | Navigator cadet | Pilot cadet | Navigator | Pilot | Navigator | Pilot |
| --- | --- | --- | --- | --- | --- | --- |
| *Aircraft* | T-6 |  | F-15 |  | F-16 |  |
| *n* | 5 | 10 | 3 | 4 | 10 | 13 |
| *Percentages out of entire sample* | 11% | 22% | 7% | 9% | 22% | 29% |
| *Average age* | 19.5 | 19.5 | 38.2 | 40.4 | 36.0 | 38.1 |
| *Number of flight profiles* | 20 | 42 | 20 | 28 | 81 | 92 |

Active aircrew members (n=30, average age = 37.74±5.46) performed all 7 profiles. Some participants performed BHA and DT twice, and one did not complete the entire procedure (as elaborated above in *Table 2*), summing to 221 sorties. Flying cadets (n=15) performed only 4 of those profiles (two of each type: the visual illusions BHA and GLS, and the vestibular illusions GLS and Coriolis) and two pilot cadets performed the BHA profile twice, summing to 62 profiles. Overall, 283 profiles were used in analyses. The flight experiment of the aircrew members was deduced from their age, as they started pilot training at 18, the flight experience of the flying cadets when undergoing the SD training during the course is 1.5 years.

Out of the 283 profiles, 136 profiles contained an SD response, i.e. it managed to induce SD. One flying cadet did not experience SD in any sorties, 2 experienced SD in 1 sortie, the rest of the cadets experienced SD either 2 or 3 of the 4 sorties. All aircrew members experienced SD during between 1 to 5 of the 7 sorties, with the majority experiencing SD in 3 sorties. The percentages of flight profiles containing an SD response was calculated per group (*Figure 6*).

**Figure 6.**
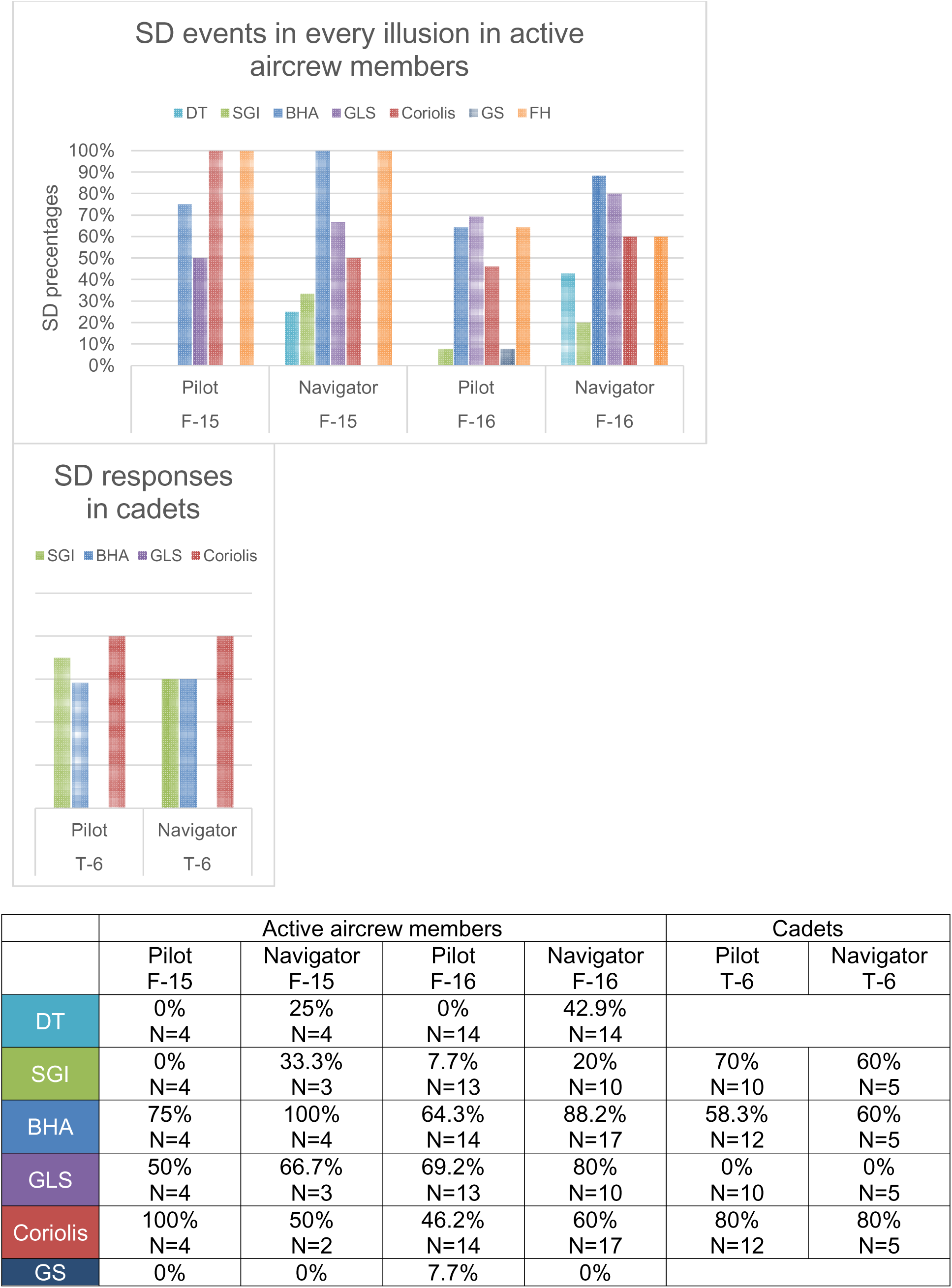
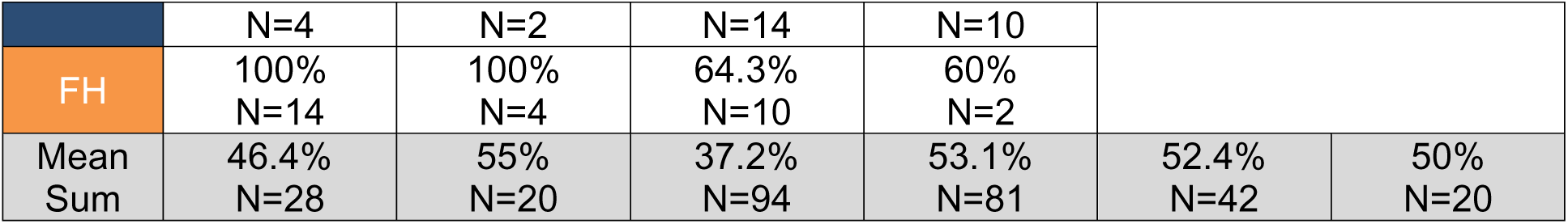
percentages for SD responses by aircraft and position. The percentage of SD responses and the total number of flight profiles for each category is shown at the table below the graph.

**Figure 7.**
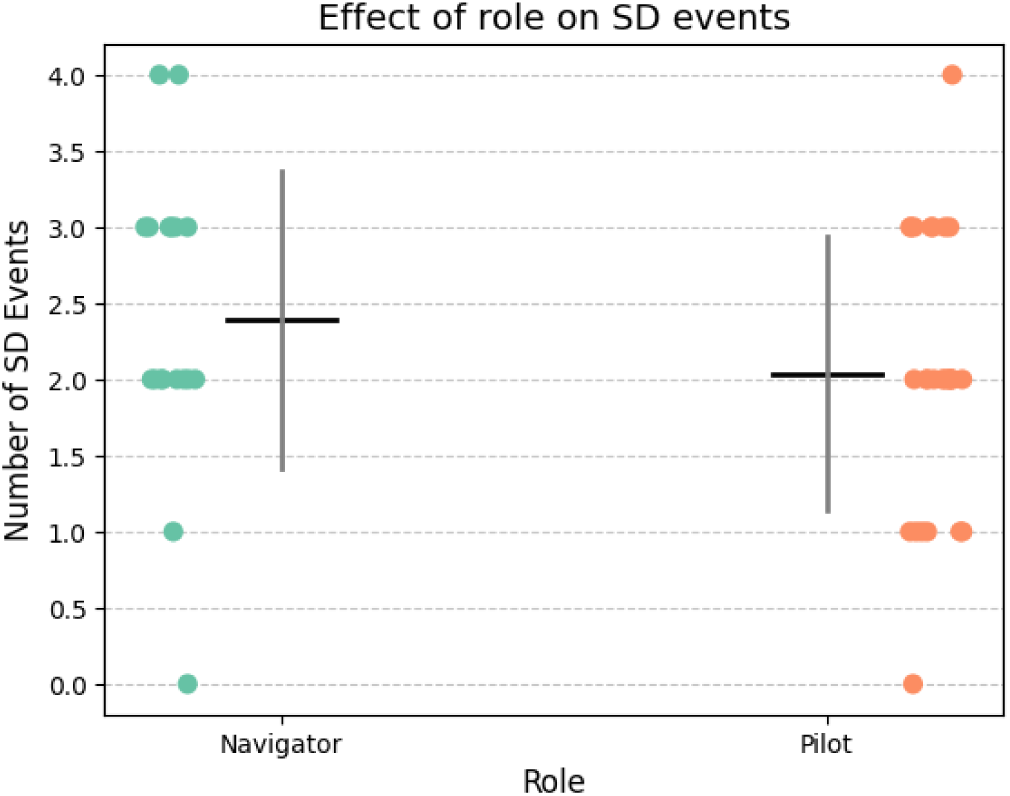
Effect of role on SD events.

We used logistic regression to examine the effects of experience and role within the aircraft on the probability of experiencing SD, while accounting for participant-level random effects. When tested separately, experience showed no significant effects— neither when modeled as a continuous variable (years of experience) nor as a binary variable (cadets vs. active aircrew members). A marginal effect was found for role, with navigators experiencing more SD responses (p = 0.06). No significant interaction between experience and role was observed.

### Eye-movement patterns

The eye gaze throughout the flight was analyzed to create heatmaps (see *Figure 8*).

**Figure 8.**
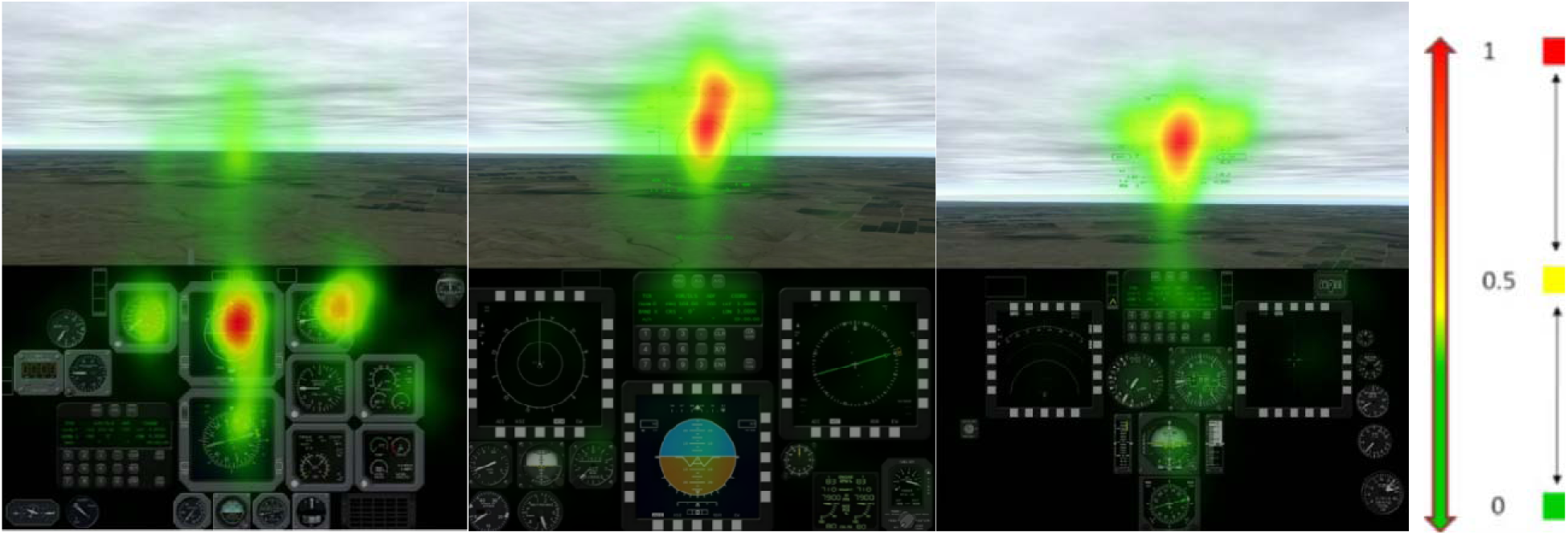
gaze heat map. Heat map of each cockpit display as calculated in the Tobii pro lab program using the Tobii I-VT (Fixation) gaze filter, from left to right: T-6, F-15, F-16. On the left there is a colorbar presenting a linear interpolation of number of fixations in each area.

The heat map showed that T-6 pilots and navigators, visit the instruments more than the external view, particularly on the ball instrument that indicates the roll and pitch degree, while F-15 and F-16 pilots visited the HUD more. The percentage of the intervals spent fixating on the different AOI categories is brought in *Table 3*:

**Table 3.**
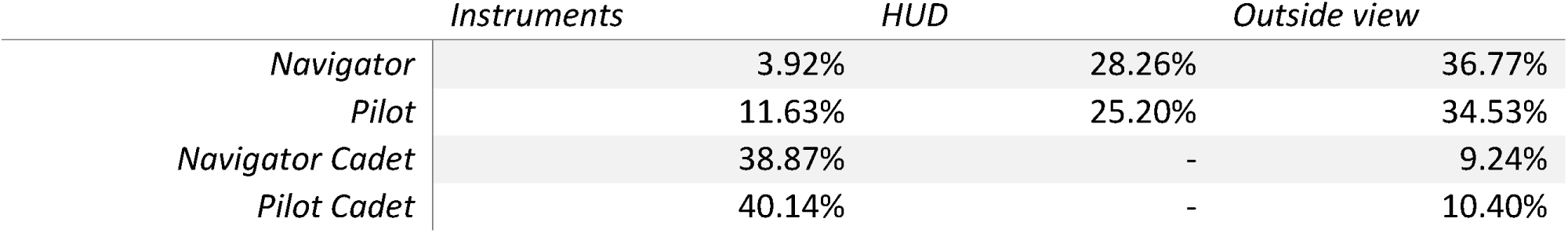
fixation time to AOIs.

|  | Instruments | HUD | Outside view |
| --- | --- | --- | --- |
| Navigator | 3.92% | 28.26% | 36.77% |
| Pilot | 11.63% | 25.20% | 34.53% |
| Navigator Cadet | 38.87% | - | 9.24% |
| Pilot Cadet | 40.14% | - | 10.40% |

### SD window analyses

To simplify and focus our analyses we categorized the four illusions experienced by all participants into visual illusions (BHA and GLS) and vestibular illusions (SGI and DT). Additionally, we grouped the Areas of Interest (AOIs) into instrument AOIs and HUD AOIs and selected five key eye movement parameters for each AOI: number of visits, number of fixations, number of saccades, total duration of visits, and total duration of fixations.

Using these data, we conducted logistic regression analyses separately for vestibular and visual illusion sorties, and separately for eye movement parameters related to instrument and HUD AOIs. Each model predicted the likelihood of an SD response based on the eye movement parameter, the role of the participant (navigator vs. pilot), and their interaction, while accounting for participant-level random effects.

After correcting for multiple comparisons, two results remained significant: within the visual illusions, longer total durations of fixations and visits on the instrument AOIs were associated with fewer SD responses (*p* < 0.001 for both; Figure 9).

**Figure 9.**
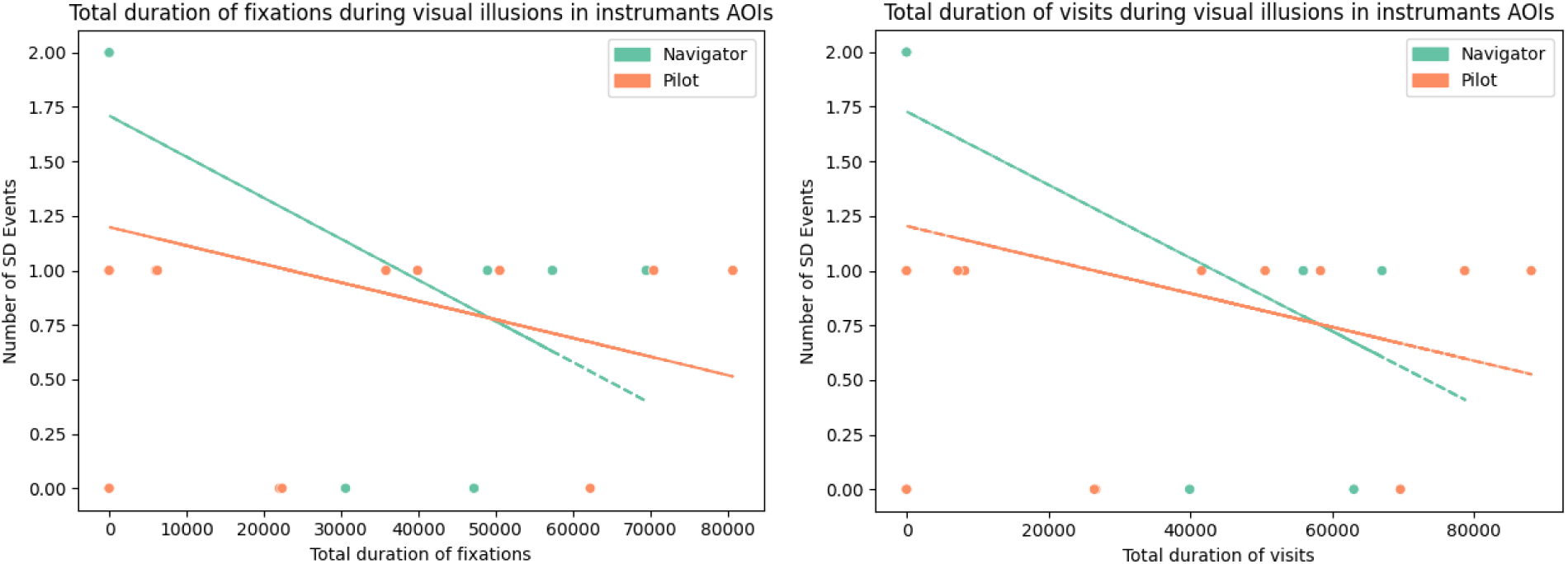
Total duration of eye tracking parameter during visual illusiond in instrumants AOIs. To further explore the interaction between illusion type (visual vs. vestibular), display type (HUD vs. instruments), and the eye tracking parameters, we ran a model testing a three-way interaction. The following effects were found (A main effect for total duration of fixations and total duration of visits, with longer durations predicting increased SD responses (p < 0.001 for both).

**Figure 10.**
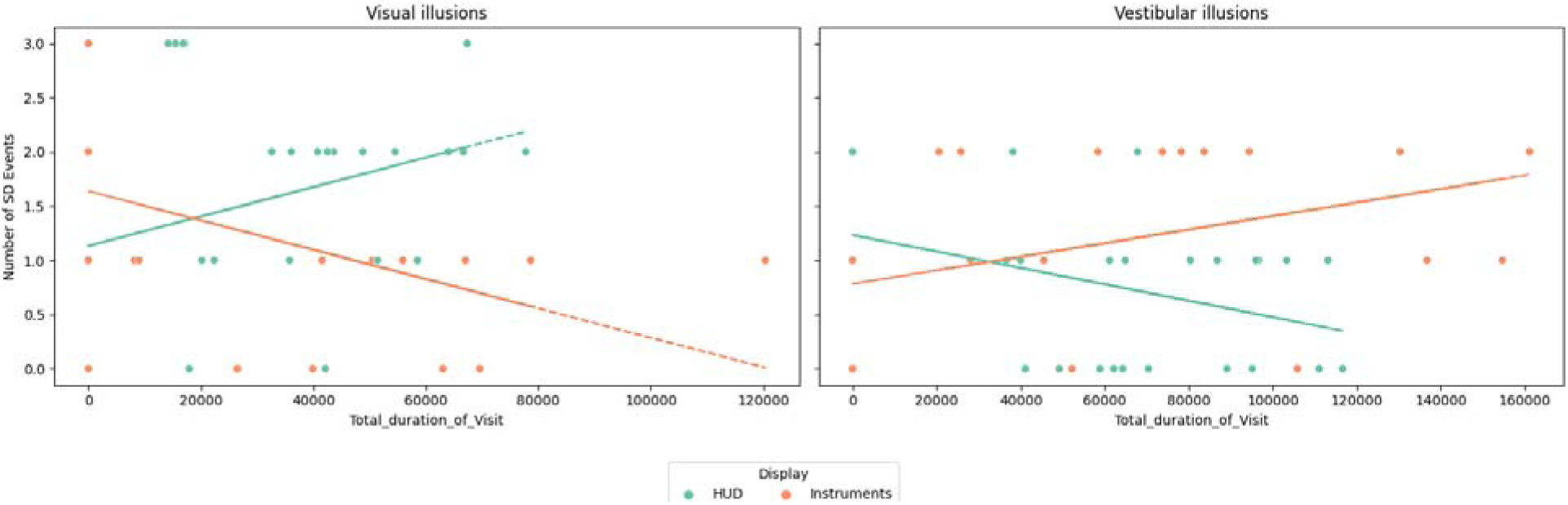
combined effect in total duration of visits.

A two-way interaction between eye tracking parameter and display type for total duration of visits, total duration of fixations, number of visits, and number of fixations: greater engagement with instrument AOIs was linked to higher SD occurrences, while greater engagement with the HUD was linked to lower SD occurrences (p < 0.001 for all).

A two-way interaction between eye tracking parameter and the illusion type for total duration of visits, total duration of fixations, and number of visits: greater engagement with the AOIs during vestibular was linked to higher SD occurrences, while in visual illusions it was linked to lower SD occurrences (p < 0.001 for all).

A two-way interaction between display type and illusion type for total duration of visits, number of visits, and number of fixations: increased HUD engagement was associated with more SD events in visual illusions but fewer in vestibular illusions, whereas instrument engagement showed the opposite pattern (p < 0.001 for all).

A triple interaction between the three arguments for total duration of visits, total duration of fixations, number of visits, and number of fixations: combining all the effects showed above (p < 0.001 for all).

Figure ):

A main effect for total duration of fixations and total duration of visits, with longer durations predicting increased SD responses (p < 0.001 for both).

A two-way interaction between eye tracking parameter and display type for total duration of visits, total duration of fixations, number of visits, and number of fixations: greater engagement with instrument AOIs was linked to higher SD occurrences, while greater engagement with the HUD was linked to lower SD occurrences (p < 0.001 for all).

A two-way interaction between eye tracking parameter and the illusion type for total duration of visits, total duration of fixations, and number of visits: greater engagement with the AOIs during vestibular was linked to higher SD occurrences, while in visual illusions it was linked to lower SD occurrences (p < 0.001 for all).

A two-way interaction between display type and illusion type for total duration of visits, number of visits, and number of fixations: increased HUD engagement was associated with more SD events in visual illusions but fewer in vestibular illusions, whereas instrument engagement showed the opposite pattern (p < 0.001 for all).

A triple interaction between the three arguments for total duration of visits, total duration of fixations, number of visits, and number of fixations: combining all the effects showed above (p < 0.001 for all).

## Discussion

In this study, we investigated eye-tracking patterns during exposure to the common illusions that cause spatial disorientation (SD) during flight both visual and vestibular. We aimed to identify which eye-tracking patterns could be related to SD avoidance.

Among 284 flight profiles performed by 30 active aircrew members and 15 flying cadets, 136 SD responses were identified across 7 flight profiles divided into **visual illusions**: Black-Hole Approach (BHA), Ground-lights and Stars (GS), and False Horizon (FH); and **vestibular illusions**: Dark Takeoff (DT), Somatogyral illusion (SGI), Coriolis, and Graveyard Spin (GS). SGI illusion was found mainly in navigators, and DT only in navigators. FH had a higher effect on F-15 aircrew than F-16, as it affected all F-15 aircrew members and only partially on F-16 aircrew members. GLS affected only active aircrew members as opposed to cadets.

Using eye-tracking we found that visual illusions had a higher probability of causing SD responses in participants who examined the instruments’ panel measurements (altitude, pitch, and ball) in greater frequencies than the HUD. In contrast, vestibular illusions showed the opposite effect of having a higher probability of causing SD responses in participants who examined the HUD in greater frequencies than the instruments’ panel measurements (altitude, pitch, and ball).

The study shows that no population avoided SD responses completely, as they affected cadets (in 50% to 52.4% of the sorties) and active aircrew members (in 37.2% to 55% of sorties), and GLS and BHA illusions even affected active pilots to a greater degree than cadets. This result is similar to a previous study that showed that SD cues affected pilots and nonpilots similarly ability to navigate the aircraft as per instructions (Bałaj et al., 2018).

In a previous study performed on transportation pilots, HUD usage increased in critical flight stages, particularly in difficult weather conditions when there was an increase in the amount of time the pilot spends looking out of the front window (Dill & Young, 2015). While the transportation pilots in that study used the HUD only 4% of the overall time, combat pilots in the current research engaged heavily on the HUD, around 90% of the overall time. This might be due to the need for high surrounding awareness as they must constantly assess external threats thus requiring a dynamic use of external visual information (Zhang et al., 2014, Martins, 2016). This effect might have occurred because the HUD allowed them to maintain visual contact with the external environment while monitoring critical flight data (Cummings et al., 2016, Johnson et al., 2006), thus a HUD usage is beneficial to reduce SD responds and is generally more relayed upon.

Our results across the two types of illusions presented to the aircrew members – visual vs. vestibular, suggest that the HUD is irrelevant in visual illusions and those who visited it numerous times for longer durations succeeded less in the event of instruments shut down (GLS).

In the vestibular illusions SGI, an increased time spent examining the HUD and a decreased time examining the instruments panel was related to better SD avoidance. This finding is supported by a former comparison of how experts’ and novices’ movement patterns change between easy and difficult routes during flight (Yang et al., 2013). For the easy routes, experts spent less time scanning out the window (OTW) and had shorter OTW dwellings. In contrast, for the difficult routes, experts appeared to slow down their scan by spending as much time scanning out the window as the novices while also having fewer Map fixations and shorter OTW dwellings. The HUD allows aircrew members to scan OTW view while examining instruments, thus allowing them to relay on the HUD in challenging flight paths.

The focus specifically directed to the HUD rather than generally examining the OTW is supported by another study that found that experienced pilots had better situational awareness performance and paid more attention to the HUD, but focused less outside the cockpit when compare with novice pilots (Yu et al., 2016). Another supporting finding to the increased need for HUD scanning to avoid SD during vestibular illusions comes from research where Coriolis was found to elevate the use of the HUD compared to other illusions and yielded a higher attentional load score (Kim et al., 2021).

### Applications

Repeated spatial training using a simulated environment has been shown to be beneficial to reduce SD, as was shown by a study conducted on neurologic patients (Kober et al., 2013), as well as in pilots (Landman et al., 2018, Qiu et al., 2023). It was found that training eye movements based on knowledge acquired about which eye movement type was the most efficient for a specific task could improve performance in said task (Shapiro & Raymond, 1989). Thus, implementing the knowledge and training based on the current results can potentially improve performance in flight profiles with high SD risk.

Another study (Peysakhovich et al., 2018) highlighted the need for real-time alert systems based on differences in eye-movement patterns associated with spatial disorientation (SD). Their study demonstrated that such systems, when installed in aircraft during real flights, can help mitigate SD-related incidents by providing timely alerts and reducing the likelihood of fatal responses.

### Limitations

Although the procedure and sample size were pre-determined in the ethics protocols, the study was exploratory and not pre-registered, leading to multiple comparisons, which hinders our knowledge on replicability of our results. Using the knowledge gained in this study, future research could create a more accurate procedure to induce stronger effects and means to address them.

### Conclusions

This study highlighted the role of targeted eye-movement behaviors underlying SD. Our results suggest that dynamic, broad scanning across instruments could reduce SD experienced in various flight illusions. Specifically, we found that flight profiles involving limited visual cues and thus a potential for visual illusions require higher attention to the HUD to avoid visually induced SD. This is opposed to prolonged turns, that induce a higher potential for vestibular illusions, required greater attention to the instruments panel to avoid SD. We also found that SD risk remains throughout all experience levels from flight cadets to experienced pilots. Training that involves lessons from this study regarding the importance of different instruments in different flight situations could improve SD resilience. Further research should broaden the findings beyond the scope of combat pilots only. In addition, examining whether training using the knowledge accumulated in the study could potentially improve SD resilience.

## Key points

- **SD Responses**: 136 SD responses across 7 flight profiles from 45 subjects; specific effects varied by role and aircraft type.
- **Illusions and Eye-Movement**: Increased instruments focus helped mitigate SD risk in visual illusions (DT, GLS, FH). Increased HUD focus improved performance vestibular illusions (SGI, Coriolis):
- **HUD Usage**: Combat pilots rely heavily on HUD (90% usage). HUD is effective to mitigate SD during vestibular illusions but less so during visual illusions.
- **Experience and SD**: Experience does not prevent SD; active pilots can be as affected as cadets.
- **Applications**: Simulated training and efficient eye-movement strategies could reduce SD. Real-time alert systems may help mitigate SD during flights.
- **Conclusion**: Training focused on tailored eye-movement and instrument use could enhance SD resilience.

## Biographies

1. Maya Harel

a. Current affiliation

i. The Israeli Air Force Aeromedical Center, Tel-Hashomer, Ramat Gan, Israel
ii. Segol School of Neuroscience, Tel Aviv University, Tel Aviv, Israel
b. Degree – M.Sc. in Neuroscience, obtained in 2025 from Tel Aviv University.
2. Idan Nakdimon

a. Current affiliation

i. The Israeli Air Force Aeromedical Center, Tel-Hashomer, Ramat Gan, Israel
b. Degree – M.Sc. in Medicine Sciences, obtained in 2019 from Tel Aviv University.
3. Oded Ben-Ari

a. Current affiliation

i. Department of Military Medicine, Faculty of Medicine, The Hebrew University of Jerusalem, Israel
ii. The Adelson School of Medicine, Ariel University, Ariel, Israel
b. Degree – MD obtained in 2009 from Tel Aviv University
4. Tom Schonberg

a. Current affiliation

i. School of Neurobiology, Biochemistry and Biophysics, The George S. Wise Faculty of Life Sciences and Sagol School of Neuroscience, Tel Aviv University, Israel
b. Degree – PhD (Department of Neurobiology and Sagol School of Neuroscience), obtained in 2010 from Tel Aviv University.

