## Supplemental Materials for "Eye movement patterns under exposure to spatial disorientation illusions during simulated flight"

Supplementary Material

### Methods

For each illusion the training profile, the SD window, criteria for identifying SD response, and a visual example of the profile timeline are presented

| **Time** | **Event** | **Explanation** |
| --- | --- | --- |
| **Vestibular illusions** | | |
| **Dark Takeoff (DT) – only for aircrew members** | | |
| 0s | A dark night,  Aircraft waiting on takeoff strip |  |
| 4s | ATC message: climb to altitude 4000ft and maintain height |  |
| Condition 1 | Is airspeed > 3 knots (1.54 m/s) |  |
| +0.01s *(from time when condition 1 is met)* | Pitch = 5°, 1.2 °/s, 2 °/s^2^, 8.10s | Adjusting pitch to simulate forward acceleration |
| Condition 2 | Is Radio Altitude > 5 ft (1.52m) |  |
| +0.01s *(from time when condition 2 is met)* | Pitch = 23°, 1°/s, 0.5°/s^2^, 20 s | Increasing pitch to combine effect of pitch and acceleration |
| +4s | HUD turns blank |  |
| SD window time,  SD response: Altitude <=1000 m & Pitch Position <= 0°  Indicates a decent when the aircraft is still too low | | |
| +19s | HUD returns to normal |  |
| Visual timeline | | |

Altitude, pitch and velocity timeline

| Time [s] | Altitude [m] | Pitch [deg] | Airspeed [m/s] |
| --- | --- | --- | --- |
| 0 | 1.451 | 0 | 0 |
| 0.0079987 | 1.451 | 0 | 0 |
| 0.0159972 | 1.45092 | 0 | 0.0259184 |
| 0.0239959 | 1.45062 | 0 | 0.0509851 |
| 0.0319943 | 1.45014 | 0 | 0.0713041 |
| 0.039993 | 1.44951 | 0 | 0.0875855 |
| 0.0479915 | 1.44876 | 0 | 0.100439 |
| 0.0559902 | 1.44792 | 0 | 0.110388 |
| 0.0639889 | 1.44701 | 0 | 0.117881 |
| 0.0719874 | 1.44605 | 0 | 0.123302 |
| 0.0799861 | 1.44505 | 0 | 0.126982 |
| 0.0879846 | 1.44402 | 0 | 0.129203 |
| 0.0959833 | 1.44299 | 0 | 0.130206 |
| 0.103982 | 1.44194 | 0 | 0.130198 |
| 0.11198 | 1.44091 | 0 | 0.129355 |
| 0.119979 | 1.43988 | 0 | 0.127828 |
| 0.127978 | 1.43886 | 0 | 0.125745 |
| 0.135976 | 1.43786 | 0 | 0.123215 |
| 0.143975 | 1.43689 | 0 | 0.120332 |
| 0.151973 | 1.43594 | 0 | 0.117174 |
| 0.159972 | 1.43501 | 0 | 0.113807 |
| 0.167971 | 1.43411 | 0 | 0.11029 |
| 0.175969 | 1.43324 | 0 | 0.106668 |
| 0.183968 | 1.4324 | 0 | 0.102982 |
| 0.191967 | 1.43159 | 0 | 0.0992652 |
| 0.199965 | 1.43081 | 0 | 0.0955458 |
| 0.207964 | 1.43006 | 0 | 0.0918467 |
| 0.215962 | 1.42934 | 0 | 0.0881869 |
| 0.223961 | 1.42865 | 0 | 0.0845822 |
| 0.231959 | 1.42798 | 0 | 0.0810449 |
| 0.239958 | 1.42735 | 0 | 0.0775854 |
| 0.247957 | 1.42674 | 0 | 0.0742114 |
| 0.255955 | 1.42616 | 0 | 0.0709291 |
| 0.263954 | 1.4256 | 0 | 0.0677432 |
| 0.271952 | 1.42507 | 0 | 0.0646569 |
| 0.279951 | 1.42457 | 0 | 0.0616723 |
| 0.28795 | 1.42409 | 0 | 0.0587908 |
| 0.295948 | 1.42363 | 0 | 0.0560128 |
| 0.303947 | 1.42319 | 0 | 0.0533383 |
| 0.311945 | 1.42277 | 0 | 0.0507666 |
| 0.319944 | 1.42238 | 0 | 0.0482966 |
| 0.327943 | 1.422 | 0 | 0.0459268 |
| 0.335941 | 1.42164 | 0 | 0.0436555 |
| 0.34394 | 1.4213 | 0 | 0.0414809 |
| 0.351938 | 1.42098 | 0 | 0.0394008 |
| 0.359937 | 1.42068 | 0 | 0.037413 |
| 0.367936 | 1.42039 | 0 | 0.0355152 |
| 0.375934 | 1.42011 | 0 | 0.033705 |
| 0.383933 | 1.41985 | 0 | 0.0319801 |
| 0.391932 | 1.4196 | 0 | 0.030338 |
| 0.39993 | 1.41937 | 0 | 0.0287764 |
| 0.407929 | 1.41915 | 0 | 0.0272929 |
| 0.415927 | 1.41894 | 0 | 0.0258852 |
| 0.423926 | 1.41874 | 0 | 0.0245511 |
| 0.431924 | 1.41856 | 0 | 0.0232884 |
| 0.439923 | 1.41838 | 0 | 0.0220951 |
| 0.447922 | 1.41821 | 0 | 0.020969 |
| 0.45592 | 1.41806 | 0 | 0.0199083 |
| 0.463919 | 1.41791 | 0 | 0.018911 |
| 0.471917 | 1.41777 | 0 | 0.0179754 |
| 0.479916 | 1.41764 | 0 | 0.0170998 |
| 0.487915 | 1.41751 | 0 | 0.0162824 |
| 0.495913 | 1.41739 | 0 | 0.0155216 |
| 0.503912 | 1.41728 | -0.00296191 | 0.0148157 |
| 0.51191 | 1.41718 | -0.00306872 | 0.0141629 |
| 0.519909 | 1.41708 | -0.00317553 | 0.0135613 |
| 0.527908 | 1.41699 | -0.00328234 | 0.0130092 |
| 0.535906 | 1.41691 | -0.00342476 | 0.0125047 |
| 0.543905 | 1.41682 | -0.00356717 | 0.0120459 |
| 0.551903 | 1.41675 | -0.00369179 | 0.011631 |
| 0.559902 | 1.41668 | -0.003852 | 0.011258 |
| 0.567901 | 1.41661 | -0.00399442 | 0.0109248 |
| 0.5759 | 1.41655 | -0.00417244 | 0.010629 |
| 0.583899 | 1.41649 | -0.00435046 | 0.0103682 |
| 0.591897 | 1.41643 | -0.00452848 | 0.0101398 |
| 0.599896 | 1.41638 | -0.0047065 | 0.00994078 |
| 0.607895 | 1.41633 | -0.00488451 | 0.00976783 |
| 0.615893 | 1.41629 | -0.00506253 | 0.00961753 |
| 0.623892 | 1.41625 | -0.00525835 | 0.00948622 |
| 0.63189 | 1.41621 | -0.00541857 | 0.00937025 |
| 0.639889 | 1.41617 | -0.00561439 | 0.00926624 |
| 0.647888 | 1.41614 | -0.00577461 | 0.00917126 |
| 0.655886 | 1.41611 | -0.00597043 | 0.00908276 |
| 0.663885 | 1.41608 | -0.00614845 | 0.00899877 |
| 0.671883 | 1.41605 | -0.00634427 | 0.00891761 |
| 0.679882 | 1.41602 | -0.00650449 | 0.00883792 |
| 0.687881 | 1.416 | -0.00668251 | 0.00875847 |
| 0.695879 | 1.41598 | -0.00684273 | 0.00867819 |
| 0.703878 | 1.41596 | -0.00700294 | 0.00859616 |
| 0.711876 | 1.41594 | -0.00716316 | 0.00851183 |
| 0.719875 | 1.41592 | -0.00732338 | 0.00842485 |
| 0.727874 | 1.4159 | -0.00744799 | 0.00833508 |
| 0.735872 | 1.41589 | -0.0075726 | 0.00824258 |
| 0.743871 | 1.41587 | -0.00769722 | 0.00814753 |
| 0.751869 | 1.41586 | -0.00782183 | 0.00805014 |
| 0.759868 | 1.41585 | -0.00796425 | 0.00795049 |
| 0.767867 | 1.41584 | -0.00803545 | 0.00784861 |
| 0.775865 | 1.41583 | -0.00819567 | 0.00774456 |
| 0.783864 | 1.41582 | -0.00828468 | 0.00763848 |
| 0.791862 | 1.41581 | -0.00840929 | 0.00753061 |
| 0.799861 | 1.4158 | -0.00858731 | 0.00742122 |
| 0.80786 | 1.4158 | -0.00865852 | 0.00731057 |
| 0.815858 | 1.41579 | -0.00881874 | 0.00719893 |
| 0.823857 | 1.41578 | -0.00894335 | 0.0070866 |
| 0.831855 | 1.41578 | -0.00906797 | 0.00697389 |
| 0.839854 | 1.41577 | -0.00919258 | 0.006861 |
| 0.847853 | 1.41577 | -0.00931719 | 0.00674799 |
| 0.855851 | 1.41577 | -0.00945961 | 0.00663493 |
| 0.86385 | 1.41576 | -0.00958422 | 0.00652182 |
| 0.871848 | 1.41576 | -0.00970883 | 0.0064087 |
| 0.879847 | 1.41576 | -0.00983345 | 0.00629566 |
| 0.887846 | 1.41575 | -0.00999367 | 0.00618286 |
| 0.895844 | 1.41575 | -0.0101183 | 0.00607046 |
| 0.903843 | 1.41575 | -0.0102429 | 0.00595859 |
| 0.911841 | 1.41575 | -0.0103675 | 0.00584735 |
| 0.91984 | 1.41575 | -0.0105277 | 0.00573686 |
| 0.927839 | 1.41574 | -0.0106167 | 0.00562719 |
| 0.935837 | 1.41574 | -0.0107948 | 0.0055184 |
| 0.943836 | 1.41574 | -0.0109016 | 0.00541052 |
| 0.951834 | 1.41574 | -0.0110262 | 0.00530357 |
| 0.959833 | 1.41574 | -0.0111686 | 0.00519759 |
| 0.967832 | 1.41574 | -0.0112932 | 0.00509268 |
| 0.97583 | 1.41574 | -0.0114178 | 0.00498897 |
| 0.983829 | 1.41574 | -0.0115246 | 0.00488662 |
| 0.991827 | 1.41574 | -0.0116492 | 0.00478573 |
| 0.999826 | 1.41573 | -0.0118095 | 0.0046864 |
| 1.00782 | 1.41573 | -0.0118985 | 0.00458872 |
| 1.01582 | 1.41573 | -0.0120231 | 0.00449278 |
| 1.02382 | 1.41573 | -0.0121299 | 0.00439867 |
| 1.03182 | 1.41573 | -0.0122189 | 0.00430642 |
| 1.03982 | 1.41573 | -0.0123435 | 0.00421604 |
| 1.04782 | 1.41573 | -0.0124325 | 0.00412756 |
| 1.05582 | 1.41573 | -0.0125037 | 0.00404099 |
| 1.06381 | 1.41573 | -0.0126283 | 0.00395634 |
| 1.07181 | 1.41573 | -0.0126996 | 0.00387363 |
| 1.07981 | 1.41573 | -0.0127886 | 0.00379284 |
| 1.08781 | 1.41573 | -0.0128598 | 0.00371395 |
| 1.09581 | 1.41572 | -0.0129666 | 0.00363696 |
| 1.10381 | 1.41572 | -0.0130378 | 0.00356183 |
| 1.11181 | 1.41572 | -0.013109 | 0.00348852 |
| 1.11981 | 1.41572 | -0.0131802 | 0.00341701 |
| 1.1278 | 1.41572 | -0.0132514 | 0.00334725 |
| 1.1358 | 1.41572 | -0.0133404 | 0.00327922 |
| 1.1438 | 1.41572 | -0.0134116 | 0.00321289 |
| 1.1518 | 1.41572 | -0.0134828 | 0.00314827 |
| 1.1598 | 1.41572 | -0.0136075 | 0.00308535 |
| 1.1678 | 1.41571 | -0.0136787 | 0.00302415 |
| 1.1758 | 1.41571 | -0.0137677 | 0.00296471 |
| 1.18379 | 1.41571 | -0.0138389 | 0.00290703 |
| 1.19179 | 1.41571 | -0.0138923 | 0.00285104 |
| 1.19979 | 1.41571 | -0.0139635 | 0.0027967 |
| 1.20779 | 1.41571 | -0.0140525 | 0.00274391 |
| 1.21579 | 1.41571 | -0.0141237 | 0.00269263 |
| 1.22379 | 1.4157 | -0.0141949 | 0.00264281 |
| 1.23179 | 1.4157 | -0.0142305 | 0.00259443 |
| 1.23978 | 1.4157 | -0.0143195 | 0.00254747 |
| 1.24778 | 1.4157 | -0.0143729 | 0.00250194 |
| 1.25578 | 1.4157 | -0.0144085 | 0.00245781 |
| 1.26378 | 1.4157 | -0.0144797 | 0.00241508 |
| 1.27178 | 1.41569 | -0.0145154 | 0.00237374 |
| 1.27978 | 1.41569 | -0.0145866 | 0.00233378 |
| 1.28778 | 1.41569 | -0.0146222 | 0.00229521 |
| 1.29577 | 1.41569 | -0.0146934 | 0.00225803 |
| 1.30377 | 1.41569 | -0.014729 | 0.00222224 |
| 1.31177 | 1.41568 | -0.0147646 | 0.0021878 |
| 1.31977 | 1.41568 | -0.0148358 | 0.00215468 |
| 1.32777 | 1.41568 | -0.0148536 | 0.00212287 |
| 1.33577 | 1.41568 | -0.0148892 | 0.00209232 |
| 1.34377 | 1.41568 | -0.0149248 | 0.00206303 |
| 1.35176 | 1.41568 | -0.014996 | 0.00203497 |
| 1.35976 | 1.41567 | -0.0150316 | 0.00200815 |
| 1.36776 | 1.41567 | -0.0150672 | 0.00198256 |
| 1.37576 | 1.41567 | -0.015085 | 0.00195817 |
| 1.38376 | 1.41567 | -0.0151206 | 0.00193497 |
| 1.39176 | 1.41567 | -0.0151562 | 0.00191293 |
| 1.39976 | 1.41566 | -0.0151918 | 0.00189199 |
| 1.40775 | 1.41566 | -0.0152274 | 0.00187215 |
| 1.41575 | 1.41566 | -0.015263 | 0.00185337 |
| 1.42375 | 1.41566 | -0.0152808 | 0.00183562 |
| 1.43175 | 1.41566 | -0.015263 | 0.00181888 |
| 1.43975 | 1.41566 | -0.0152986 | 0.00180309 |
| 1.44775 | 1.41565 | -0.0153342 | 0.00178824 |
| 1.45575 | 1.41565 | -0.0153698 | 0.00177427 |
| 1.46374 | 1.41565 | -0.015352 | 0.00176117 |
| 1.47174 | 1.41565 | -0.0153698 | 0.00174891 |
| 1.47974 | 1.41565 | -0.0153698 | 0.00173746 |
| 1.48774 | 1.41565 | -0.0153876 | 0.00172682 |
| 1.49574 | 1.41564 | -0.0153876 | 0.00171695 |
| 1.50374 | 1.41564 | -0.0153876 | 0.00170783 |
| 1.51174 | 1.41564 | -0.0153698 | 0.00169945 |
| 1.51974 | 1.41564 | -0.0154054 | 0.00169179 |
| 1.52773 | 1.41564 | -0.0153876 | 0.00168481 |
| 1.53573 | 1.41564 | -0.0154054 | 0.0016785 |
| 1.54373 | 1.41564 | -0.0153876 | 0.00167282 |
| 1.55173 | 1.41563 | -0.015352 | 0.00166778 |
| 1.55973 | 1.41563 | -0.0153876 | 0.00166334 |
| 1.56773 | 1.41563 | -0.015352 | 0.00165946 |
| 1.57573 | 1.41563 | -0.0153342 | 0.00165607 |
| 1.58372 | 1.41563 | -0.0153342 | 0.00165313 |
| 1.59172 | 1.41563 | -0.0153342 | 0.00165063 |

| **Somatogyral Illusion (SGI)** | | |
| --- | --- | --- |
| 0s | A dark night, Aircraft flying horizontally and leveled, Aircraft altitude = 7000 ft, heading point = 25° |  |
| 20s | ATC message 1: turn right 45° and maintain aircraft tilt until further instructions |  |
| Condition | Roll angle not between -5° and 5° | Roll motion begins |
| +22s | City lights turned off | Reduces visual cues |
| +39s | Cumulus clouds appear (8/8 coverage, between 3000 and 6000 ft) |  |
| +40s | ATC message 2: return to straight and leveled flight |  |
| SD window time,  SD response: Roll Position > 0° & Roll Velocity >= 3 °/s  Indicates a motion towards original roll | | |
| +60s |  |  |
| Visual timeline | | |

Roll and heading timeline

| Time [s] | Roll [deg] | Heading [deg] |
| --- | --- | --- |
| 0 | 1.42011e-13 | 25.2728 |
| 0.0079987 | 1.42011e-13 | 25.2728 |
| 0.0159974 | 2.70499e-08 | 25.2728 |
| 0.0239961 | 5.79656e-08 | 25.2728 |
| 0.0319951 | 8.88829e-08 | 25.2728 |
| 0.0399938 | 1.19802e-07 | 25.2728 |
| 0.0479925 | 1.50722e-07 | 25.2728 |
| 0.0559912 | 1.81644e-07 | 25.2728 |
| 0.0639899 | 2.12568e-07 | 25.2728 |
| 0.0719883 | 2.43493e-07 | 25.2728 |
| 0.079987 | 2.74419e-07 | 25.2728 |
| 0.0879855 | 3.05346e-07 | 25.2728 |
| 0.0959842 | 3.36275e-07 | 25.2728 |
| 0.103983 | 3.67205e-07 | 25.2728 |
| 0.111981 | 3.98135e-07 | 25.2728 |
| 0.11998 | 4.29067e-07 | 25.2728 |
| 0.127979 | 4.59999e-07 | 25.2728 |
| 0.135977 | 4.90933e-07 | 25.2728 |
| 0.143976 | 5.21867e-07 | 25.2728 |
| 0.151974 | 5.52802e-07 | 25.2728 |
| 0.159973 | 5.83738e-07 | 25.2728 |
| 0.167972 | 6.14675e-07 | 25.2728 |
| 0.17597 | 6.45613e-07 | 25.2728 |
| 0.183969 | 6.76552e-07 | 25.2728 |
| 0.191967 | 7.07491e-07 | 25.2728 |
| 0.199966 | 7.38431e-07 | 25.2728 |
| 0.207965 | 7.69372e-07 | 25.2728 |
| 0.215963 | 8.00314e-07 | 25.2728 |
| 0.223962 | 8.31256e-07 | 25.2728 |
| 0.23196 | 8.622e-07 | 25.2728 |
| 0.239959 | 8.93144e-07 | 25.2728 |
| 0.247958 | 9.24089e-07 | 25.2728 |
| 0.255956 | 9.55035e-07 | 25.2728 |
| 0.263955 | 9.85982e-07 | 25.2728 |
| 0.271953 | 1.01693e-06 | 25.2728 |
| 0.279952 | 1.04788e-06 | 25.2729 |
| 0.287951 | 1.07883e-06 | 25.2729 |
| 0.295949 | 1.10978e-06 | 25.2729 |
| 0.303948 | 1.14073e-06 | 25.2729 |
| 0.311946 | 1.17168e-06 | 25.2729 |
| 0.319945 | 1.20263e-06 | 25.2729 |
| 0.327944 | 1.23358e-06 | 25.2729 |
| 0.335942 | 1.26454e-06 | 25.2729 |
| 0.343941 | 1.29549e-06 | 25.2729 |
| 0.351939 | 1.32645e-06 | 25.2729 |
| 0.359938 | 1.3574e-06 | 25.2729 |
| 0.367937 | 1.38836e-06 | 25.2729 |
| 0.375935 | 1.41931e-06 | 25.2729 |
| 0.383934 | 1.45027e-06 | 25.2729 |
| 0.391932 | 1.48123e-06 | 25.2729 |
| 0.399931 | 1.51219e-06 | 25.2729 |
| 0.40793 | 1.54315e-06 | 25.2729 |
| 0.415928 | 1.57411e-06 | 25.2729 |
| 0.423927 | 1.60507e-06 | 25.2729 |
| 0.431925 | 1.63603e-06 | 25.2729 |
| 0.439924 | 1.66699e-06 | 25.2729 |
| 0.447923 | 1.69795e-06 | 25.2729 |
| 0.455921 | 1.72892e-06 | 25.2729 |
| 0.46392 | 1.75988e-06 | 25.2729 |
| 0.471918 | 1.79084e-06 | 25.2729 |
| 0.479917 | 1.82181e-06 | 25.2729 |
| 0.487916 | 1.85277e-06 | 25.2729 |
| 0.495914 | 1.88374e-06 | 25.2729 |
| 0.503913 | 1.9147e-06 | 25.2729 |
| 0.511911 | 1.94567e-06 | 25.2729 |
| 0.51991 | 1.97664e-06 | 25.2729 |
| 0.527909 | 2.0076e-06 | 25.2729 |
| 0.535907 | 2.03857e-06 | 25.2729 |
| 0.543906 | 2.06954e-06 | 25.2729 |
| 0.551904 | 2.10051e-06 | 25.273 |
| 0.559903 | 2.13148e-06 | 25.273 |
| 0.567902 | 2.16245e-06 | 25.273 |
| 0.5759 | 2.19342e-06 | 25.273 |
| 0.583899 | 2.22439e-06 | 25.273 |
| 0.591897 | 2.25536e-06 | 25.273 |
| 0.599896 | 2.28633e-06 | 25.273 |
| 0.607895 | 2.3173e-06 | 25.273 |
| 0.615893 | 2.34827e-06 | 25.273 |
| 0.623892 | 2.37924e-06 | 25.273 |
| 0.63189 | 2.41022e-06 | 25.273 |
| 0.639889 | 2.44119e-06 | 25.273 |
| 0.647888 | 2.47217e-06 | 25.273 |
| 0.655886 | 2.50314e-06 | 25.273 |
| 0.663885 | 2.53411e-06 | 25.273 |
| 0.671883 | 2.56509e-06 | 25.273 |
| 0.679882 | 2.59606e-06 | 25.273 |
| 0.687881 | 2.62704e-06 | 25.273 |
| 0.695879 | 2.65802e-06 | 25.273 |
| 0.703878 | 2.68899e-06 | 25.273 |
| 0.711876 | 2.71997e-06 | 25.273 |
| 0.719875 | 2.75095e-06 | 25.273 |
| 0.727874 | 2.78192e-06 | 25.273 |
| 0.735872 | 2.8129e-06 | 25.273 |
| 0.743871 | 2.84388e-06 | 25.273 |
| 0.751869 | 2.87486e-06 | 25.273 |
| 0.759868 | 2.90584e-06 | 25.273 |
| 0.767867 | 2.93682e-06 | 25.273 |
| 0.775865 | 2.9678e-06 | 25.273 |
| 0.783864 | 2.99878e-06 | 25.273 |
| 0.791862 | 3.02976e-06 | 25.273 |
| 0.799861 | 3.06074e-06 | 25.273 |
| 0.80786 | 3.09172e-06 | 25.273 |
| 0.815858 | 3.12271e-06 | 25.2731 |
| 0.823857 | 3.15369e-06 | 25.2731 |
| 0.831855 | 3.18467e-06 | 25.2731 |
| 0.839854 | 3.21565e-06 | 25.2731 |
| 0.847853 | 3.24664e-06 | 25.2731 |
| 0.855851 | 3.27762e-06 | 25.2731 |
| 0.86385 | 3.3086e-06 | 25.2731 |
| 0.871848 | 3.33959e-06 | 25.2731 |
| 0.879847 | 3.37057e-06 | 25.2731 |
| 0.887846 | 3.40156e-06 | 25.2731 |
| 0.895844 | 3.43255e-06 | 25.2731 |
| 0.903843 | 3.46353e-06 | 25.2731 |
| 0.911841 | 3.49452e-06 | 25.2731 |
| 0.91984 | 3.52551e-06 | 25.2731 |
| 0.927839 | 3.55649e-06 | 25.2731 |
| 0.935837 | 3.58748e-06 | 25.2731 |
| 0.943836 | 3.61847e-06 | 25.2731 |
| 0.951834 | 3.64946e-06 | 25.2731 |
| 0.959833 | 3.68045e-06 | 25.2731 |
| 0.967832 | 3.71144e-06 | 25.2731 |
| 0.97583 | 3.74243e-06 | 25.2731 |
| 0.983829 | 3.77342e-06 | 25.2731 |
| 0.991827 | 3.80441e-06 | 25.2731 |
| 0.999826 | 3.8354e-06 | 25.2731 |
| 1.00782 | 3.8354e-06 | 25.2731 |
| 1.01582 | 3.86639e-06 | 25.2731 |
| 1.02382 | 3.92837e-06 | 25.2731 |
| 1.03182 | 3.95937e-06 | 25.2731 |
| 1.03982 | 3.99036e-06 | 25.2731 |
| 1.04782 | 4.02135e-06 | 25.2731 |
| 1.05582 | 4.05235e-06 | 25.2731 |
| 1.06382 | 4.08334e-06 | 25.2731 |
| 1.07181 | 4.11434e-06 | 25.2731 |
| 1.07981 | 4.14533e-06 | 25.2731 |
| 1.08781 | 4.17633e-06 | 25.2732 |
| 1.09581 | 4.20733e-06 | 25.2732 |
| 1.10381 | 4.23832e-06 | 25.2732 |
| 1.11181 | 4.26932e-06 | 25.2732 |
| 1.11981 | 4.30032e-06 | 25.2732 |
| 1.1278 | 4.33132e-06 | 25.2732 |
| 1.1358 | 4.36231e-06 | 25.2732 |
| 1.1438 | 4.39331e-06 | 25.2732 |
| 1.1518 | 4.42431e-06 | 25.2732 |
| 1.1598 | 4.45531e-06 | 25.2732 |
| 1.1678 | 4.48632e-06 | 25.2732 |
| 1.1758 | 4.51732e-06 | 25.2732 |
| 1.18379 | 4.54832e-06 | 25.2732 |
| 1.19179 | 4.57932e-06 | 25.2732 |
| 1.19979 | 4.61032e-06 | 25.2732 |
| 1.20779 | 4.64133e-06 | 25.2732 |
| 1.21579 | 4.67233e-06 | 25.2732 |
| 1.22379 | 4.70334e-06 | 25.2732 |
| 1.23179 | 4.73434e-06 | 25.2732 |
| 1.23978 | 4.76535e-06 | 25.2732 |
| 1.24778 | 4.79635e-06 | 25.2732 |
| 1.25578 | 4.82736e-06 | 25.2732 |
| 1.26378 | 4.85837e-06 | 25.2732 |
| 1.27178 | 4.88938e-06 | 25.2732 |
| 1.27978 | 4.92039e-06 | 25.2732 |
| 1.28778 | 4.9514e-06 | 25.2732 |
| 1.29577 | 4.9824e-06 | 25.2732 |
| 1.30377 | 5.01342e-06 | 25.2732 |
| 1.31177 | 5.04443e-06 | 25.2732 |
| 1.31977 | 5.07544e-06 | 25.2732 |
| 1.32777 | 5.10645e-06 | 25.2732 |
| 1.33577 | 5.13746e-06 | 25.2732 |
| 1.34377 | 5.16848e-06 | 25.2732 |
| 1.35177 | 5.19949e-06 | 25.2733 |
| 1.35976 | 5.23051e-06 | 25.2733 |
| 1.36776 | 5.26152e-06 | 25.2733 |
| 1.37576 | 5.29254e-06 | 25.2733 |
| 1.38376 | 5.32356e-06 | 25.2733 |
| 1.39176 | 5.35457e-06 | 25.2733 |
| 1.39976 | 5.38559e-06 | 25.2733 |
| 1.40776 | 5.41661e-06 | 25.2733 |
| 1.41575 | 5.44763e-06 | 25.2733 |
| 1.42375 | 5.47865e-06 | 25.2733 |
| 1.43175 | 5.50967e-06 | 25.2733 |
| 1.43975 | 5.54069e-06 | 25.2733 |
| 1.44775 | 5.57172e-06 | 25.2733 |
| 1.45575 | 5.60274e-06 | 25.2733 |
| 1.46375 | 5.63377e-06 | 25.2733 |
| 1.47174 | 5.66479e-06 | 25.2733 |
| 1.47974 | 5.69582e-06 | 25.2733 |
| 1.48774 | 5.72684e-06 | 25.2733 |
| 1.49574 | 5.75787e-06 | 25.2733 |
| 1.50374 | 5.7889e-06 | 25.2733 |
| 1.51174 | 5.81993e-06 | 25.2733 |
| 1.51974 | 5.85096e-06 | 25.2733 |
| 1.52773 | 5.88199e-06 | 25.2733 |
| 1.53573 | 5.91302e-06 | 25.2733 |
| 1.54373 | 5.94405e-06 | 25.2733 |
| 1.55173 | 5.97508e-06 | 25.2733 |
| 1.55973 | 6.00612e-06 | 25.2733 |
| 1.56773 | 6.03715e-06 | 25.2733 |
| 1.57573 | 6.06819e-06 | 25.2733 |
| 1.58372 | 6.09923e-06 | 25.2733 |
| 1.59172 | 6.13026e-06 | 25.2733 |

| **Coriolis** | | |
| --- | --- | --- |
| 0s | Aircraft flying horizontally and leveled, Aircraft heading towards 330°, altitude = 16,000 ft |  |
| 4s | ATC message 1: turn right towards heading point 200 | Increases heading direction |
| Condition 1 | Roll angle not between -5° and 5° | Roll motion begins |
| +0.01s | Yaw velocity = 50°/s, 3°/s², 16.67s | Stabilize turn velocity to ensure effect |
| Condition 2 | Heading > 150° |  |
| +4s | ATC message 2: continue turning towards heading point 270 | Continue turn |
| Condition 3 | Heading > 220° |  |
| +0.01s | ATC message 3: examine a button placed behind to the left of the aircrew member | Requires a backwards head movement from the aircrew member |
| SD window time,  SD response: Roll Velocity <-7°/s or > 7°/s or Pitch Velocity < -5°/s or > 5°/s  Indicates a drastic stick movement due to the roll sensation | | |
| +10s |  |  |
| Visual timeline | | |

Roll and heading timeline

| Time [s] | Roll [deg] | Heading [deg] |
| --- | --- | --- |
| 0 | -6.68e-13 | 325.277 |
| 0.00799871 | -6.68e-13 | 325.277 |
| 0.0159972 | -6.68e-13 | 325.277 |
| 0.0239959 | 2.24e-05 | 325.277 |
| 0.0319943 | 0.000849669 | 325.277 |
| 0.039993 | 0.000849669 | 325.277 |
| 0.0479918 | 0.00333057 | 325.277 |
| 0.0559902 | 0.00519612 | 325.277 |
| 0.0639889 | 0.0074332 | 325.277 |
| 0.0719874 | 0.0100005 | 325.277 |
| 0.0799861 | 0.012856 | 325.277 |
| 0.0879846 | 0.0159582 | 325.277 |
| 0.0959833 | 0.0159582 | 325.277 |
| 0.103982 | 0.019267 | 325.277 |
| 0.11198 | 0.0227457 | 325.277 |
| 0.119979 | 0.0263571 | 325.277 |
| 0.127978 | 0.0300663 | 325.277 |
| 0.135976 | 0.0338427 | 325.277 |
| 0.143975 | 0.0376591 | 325.276 |
| 0.151973 | 0.041492 | 325.276 |
| 0.159972 | 0.0453214 | 325.276 |
| 0.167971 | 0.0491313 | 325.276 |
| 0.175969 | 0.0529101 | 325.276 |
| 0.183968 | 0.0566501 | 325.276 |
| 0.191967 | 0.0603462 | 325.275 |
| 0.199965 | 0.0639959 | 325.275 |
| 0.207964 | 0.0675999 | 325.275 |
| 0.215962 | 0.0711614 | 325.275 |
| 0.223961 | 0.0746856 | 325.274 |
| 0.23196 | 0.078177 | 325.274 |
| 0.239958 | 0.0816407 | 325.274 |
| 0.247957 | 0.0850823 | 325.274 |
| 0.255955 | 0.0885092 | 325.273 |
| 0.263954 | 0.0919297 | 325.273 |
| 0.271953 | 0.0953529 | 325.273 |
| 0.279951 | 0.098787 | 325.273 |
| 0.28795 | 0.10224 | 325.272 |
| 0.295949 | 0.105721 | 325.272 |
| 0.303947 | 0.109239 | 325.272 |
| 0.311946 | 0.112811 | 325.272 |
| 0.319944 | 0.116454 | 325.271 |
| 0.327943 | 0.12019 | 325.271 |
| 0.335942 | 0.124042 | 325.271 |
| 0.34394 | 0.128029 | 325.271 |
| 0.351939 | 0.132165 | 325.27 |
| 0.359937 | 0.136457 | 325.27 |
| 0.367936 | 0.140905 | 325.27 |
| 0.375935 | 0.145504 | 325.269 |
| 0.383933 | 0.150248 | 325.269 |
| 0.391932 | 0.155126 | 325.269 |
| 0.39993 | 0.160132 | 325.269 |
| 0.407929 | 0.165261 | 325.268 |
| 0.415928 | 0.170511 | 325.268 |
| 0.423926 | 0.175882 | 325.268 |
| 0.431925 | 0.181374 | 325.267 |
| 0.439924 | 0.186982 | 325.267 |
| 0.447922 | 0.192704 | 325.267 |
| 0.455921 | 0.198531 | 325.266 |
| 0.463919 | 0.204456 | 325.266 |
| 0.471918 | 0.210468 | 325.266 |
| 0.479917 | 0.216558 | 325.265 |
| 0.487915 | 0.222717 | 325.265 |
| 0.495914 | 0.228934 | 325.265 |
| 0.503912 | 0.235203 | 325.264 |
| 0.511911 | 0.241515 | 325.264 |
| 0.51991 | 0.247868 | 325.264 |
| 0.527908 | 0.254254 | 325.263 |
| 0.535907 | 0.26067 | 325.263 |
| 0.543905 | 0.26711 | 325.262 |
| 0.551904 | 0.27357 | 325.262 |
| 0.559902 | 0.280043 | 325.262 |
| 0.567901 | 0.286528 | 325.261 |
| 0.5759 | 0.293018 | 325.261 |
| 0.583898 | 0.299513 | 325.26 |
| 0.591897 | 0.306011 | 325.26 |
| 0.599895 | 0.312509 | 325.259 |
| 0.607894 | 0.319009 | 325.259 |
| 0.615893 | 0.325508 | 325.259 |
| 0.623891 | 0.332007 | 325.258 |
| 0.63189 | 0.338506 | 325.258 |
| 0.639889 | 0.345002 | 325.257 |
| 0.647887 | 0.351492 | 325.257 |
| 0.655886 | 0.357977 | 325.257 |
| 0.663884 | 0.364457 | 325.256 |
| 0.671883 | 0.370934 | 325.256 |
| 0.679882 | 0.377412 | 325.255 |
| 0.687881 | 0.383892 | 325.255 |
| 0.695879 | 0.390377 | 325.255 |
| 0.703878 | 0.396867 | 325.254 |
| 0.711877 | 0.403363 | 325.254 |
| 0.719876 | 0.40986 | 325.253 |
| 0.727874 | 0.416355 | 325.253 |
| 0.735873 | 0.422844 | 325.253 |
| 0.743871 | 0.429324 | 325.252 |
| 0.75187 | 0.435793 | 325.252 |
| 0.759869 | 0.442249 | 325.251 |
| 0.767867 | 0.448693 | 325.251 |
| 0.775866 | 0.455128 | 325.251 |
| 0.783864 | 0.461556 | 325.25 |
| 0.791863 | 0.467979 | 325.25 |
| 0.799862 | 0.474398 | 325.25 |
| 0.80786 | 0.480815 | 325.249 |
| 0.815859 | 0.487232 | 325.249 |
| 0.823858 | 0.493648 | 325.248 |
| 0.831856 | 0.500064 | 325.248 |
| 0.839855 | 0.506482 | 325.248 |
| 0.847853 | 0.512901 | 325.247 |
| 0.855852 | 0.519318 | 325.247 |
| 0.863851 | 0.525732 | 325.247 |
| 0.871849 | 0.532139 | 325.246 |
| 0.879848 | 0.538538 | 325.246 |
| 0.887846 | 0.544928 | 325.246 |
| 0.895845 | 0.551307 | 325.246 |
| 0.903844 | 0.557677 | 325.245 |
| 0.911842 | 0.564038 | 325.245 |
| 0.919841 | 0.570395 | 325.245 |
| 0.92784 | 0.57675 | 325.244 |
| 0.935838 | 0.583105 | 325.244 |
| 0.943837 | 0.58946 | 325.244 |
| 0.951835 | 0.595816 | 325.244 |
| 0.959834 | 0.602171 | 325.243 |
| 0.967833 | 0.608525 | 325.243 |
| 0.975831 | 0.614879 | 325.243 |
| 0.98383 | 0.621236 | 325.243 |
| 0.991828 | 0.6276 | 325.242 |
| 0.999827 | 0.633973 | 325.242 |
| 1.00783 | 0.640356 | 325.242 |
| 1.01582 | 0.646748 | 325.242 |
| 1.02382 | 0.653147 | 325.241 |
| 1.03182 | 0.659551 | 325.241 |
| 1.03982 | 0.665957 | 325.241 |
| 1.04782 | 0.672361 | 325.241 |
| 1.05582 | 0.678753 | 325.24 |
| 1.06382 | 0.685122 | 325.24 |
| 1.07181 | 0.691449 | 325.24 |
| 1.07981 | 0.697709 | 325.24 |
| 1.08781 | 0.703874 | 325.24 |
| 1.09581 | 0.70991 | 325.24 |
| 1.10381 | 0.715787 | 325.239 |
| 1.11181 | 0.721474 | 325.239 |
| 1.11981 | 0.726947 | 325.239 |
| 1.1278 | 0.732192 | 325.239 |
| 1.1358 | 0.7372 | 325.239 |
| 1.1438 | 0.741974 | 325.238 |
| 1.1518 | 0.746523 | 325.238 |
| 1.1598 | 0.750863 | 325.238 |
| 1.1678 | 0.755013 | 325.238 |
| 1.1758 | 0.758997 | 325.238 |
| 1.18379 | 0.76284 | 325.238 |
| 1.19179 | 0.76657 | 325.238 |
| 1.19979 | 0.770214 | 325.238 |
| 1.20779 | 0.773797 | 325.238 |
| 1.21579 | 0.777339 | 325.238 |
| 1.22379 | 0.780856 | 325.238 |
| 1.23179 | 0.78436 | 325.238 |
| 1.23979 | 0.787859 | 325.238 |
| 1.24778 | 0.791358 | 325.238 |
| 1.25578 | 0.794858 | 325.238 |
| 1.26378 | 0.798362 | 325.238 |
| 1.27178 | 0.801869 | 325.238 |
| 1.27978 | 0.805379 | 325.238 |
| 1.28778 | 0.808891 | 325.238 |
| 1.29578 | 0.812404 | 325.238 |
| 1.30377 | 0.815916 | 325.238 |
| 1.31177 | 0.819426 | 325.238 |
| 1.31977 | 0.822931 | 325.238 |
| 1.32777 | 0.826427 | 325.238 |
| 1.33577 | 0.829911 | 325.238 |
| 1.34377 | 0.833381 | 325.239 |
| 1.35177 | 0.836835 | 325.239 |
| 1.35976 | 0.840272 | 325.239 |
| 1.36776 | 0.843692 | 325.239 |
| 1.37576 | 0.847093 | 325.239 |
| 1.38376 | 0.850476 | 325.239 |
| 1.39176 | 0.853842 | 325.239 |
| 1.39976 | 0.857193 | 325.24 |
| 1.40776 | 0.860533 | 325.24 |
| 1.41575 | 0.863866 | 325.24 |
| 1.42375 | 0.867197 | 325.24 |
| 1.43175 | 0.870529 | 325.24 |
| 1.43975 | 0.873872 | 325.24 |
| 1.44775 | 0.877234 | 325.241 |
| 1.45575 | 0.880628 | 325.241 |
| 1.46375 | 0.884064 | 325.241 |
| 1.47174 | 0.887556 | 325.241 |
| 1.47974 | 0.89111 | 325.241 |
| 1.48774 | 0.894733 | 325.242 |
| 1.49574 | 0.898424 | 325.242 |
| 1.50374 | 0.90218 | 325.242 |
| 1.51174 | 0.905995 | 325.242 |
| 1.51974 | 0.909863 | 325.242 |
| 1.52774 | 0.913778 | 325.243 |
| 1.53573 | 0.917734 | 325.243 |
| 1.54373 | 0.921726 | 325.243 |
| 1.55173 | 0.925752 | 325.243 |
| 1.55973 | 0.929803 | 325.243 |
| 1.56773 | 0.933873 | 325.244 |
| 1.57573 | 0.93795 | 325.244 |
| 1.58373 | 0.94202 | 325.244 |
| 1.59172 | 0.946069 | 325.244 |

| **Graveyard Spin (GS) – only for aircrew members** | | |
| --- | --- | --- |
| 0s | Aircraft flying horizontally and leveled, Aircraft altitude = 20,500 ft (6248.4 m) |  |
| 8s | ATC message 1: enter a spin when an instructed |  |
| 43s | ATC message 2: enter spin |  |
| Condition | Altitude < 17500 ft (5334 m) | When lowering too much, instructing to recover from the spin |
| +3s | ATC message 3: recover from spin |  |
| SD window time,  SD response: Roll Position > 0° & Roll Velocity >= 3 °/s  Indicates a motion towards original spin | | |
| Visual timeline | | |

Altitude [m]

| Time [s] | aeromodel.altitude (m) |
| --- | --- |
| 0 | 6248.4 |
| 0.00799847 | 6248.4 |
| 0.0159972 | 6249.85 |
| 0.0239959 | 6249.85 |
| 0.0319943 | 6249.85 |
| 0.039993 | 6249.85 |
| 0.0479918 | 6249.85 |
| 0.0559902 | 6249.85 |
| 0.0639889 | 6249.85 |
| 0.0719874 | 6249.85 |
| 0.0799861 | 6249.85 |
| 0.0879848 | 6249.85 |
| 0.0959833 | 6249.85 |
| 0.103982 | 6249.85 |
| 0.111981 | 6249.85 |
| 0.119979 | 6249.85 |
| 0.127978 | 6249.85 |
| 0.135977 | 6249.85 |
| 0.143975 | 6249.85 |
| 0.151974 | 6249.85 |
| 0.159972 | 6249.84 |
| 0.167971 | 6249.84 |
| 0.17597 | 6249.84 |
| 0.183968 | 6249.84 |
| 0.191967 | 6249.84 |
| 0.199965 | 6249.84 |
| 0.207964 | 6249.84 |
| 0.215963 | 6249.84 |
| 0.223961 | 6249.84 |
| 0.23196 | 6249.84 |
| 0.239959 | 6249.84 |
| 0.247957 | 6249.83 |
| 0.255956 | 6249.83 |
| 0.263954 | 6249.83 |
| 0.271953 | 6249.83 |
| 0.279952 | 6249.83 |
| 0.28795 | 6249.83 |
| 0.295949 | 6249.83 |
| 0.303947 | 6249.83 |
| 0.311946 | 6249.83 |
| 0.319945 | 6249.82 |
| 0.327943 | 6249.82 |
| 0.335942 | 6249.82 |
| 0.34394 | 6249.82 |
| 0.351939 | 6249.82 |
| 0.359938 | 6249.82 |
| 0.367936 | 6249.81 |
| 0.375935 | 6249.81 |
| 0.383934 | 6249.81 |
| 0.391932 | 6249.81 |
| 0.399931 | 6249.81 |
| 0.407929 | 6249.81 |
| 0.415928 | 6249.8 |
| 0.423927 | 6249.8 |
| 0.431925 | 6249.8 |
| 0.439924 | 6249.8 |
| 0.447922 | 6249.8 |
| 0.455921 | 6249.79 |
| 0.46392 | 6249.79 |
| 0.471918 | 6249.79 |
| 0.479917 | 6249.79 |
| 0.487916 | 6249.79 |
| 0.495915 | 6249.78 |
| 0.503913 | 6249.78 |
| 0.511912 | 6249.78 |
| 0.519911 | 6249.78 |
| 0.52791 | 6249.77 |
| 0.535908 | 6249.77 |
| 0.543907 | 6249.77 |
| 0.551905 | 6249.77 |
| 0.559904 | 6249.76 |
| 0.567903 | 6249.76 |
| 0.575901 | 6249.76 |
| 0.5839 | 6249.76 |
| 0.591898 | 6249.75 |
| 0.599897 | 6249.75 |
| 0.607896 | 6249.75 |
| 0.615894 | 6249.75 |
| 0.623893 | 6249.74 |
| 0.631891 | 6249.74 |
| 0.63989 | 6249.74 |
| 0.647889 | 6249.73 |
| 0.655887 | 6249.73 |
| 0.663886 | 6249.73 |
| 0.671885 | 6249.73 |
| 0.679883 | 6249.72 |
| 0.687882 | 6249.72 |
| 0.69588 | 6249.72 |
| 0.703879 | 6249.71 |
| 0.711878 | 6249.71 |
| 0.719876 | 6249.71 |
| 0.727875 | 6249.7 |
| 0.735873 | 6249.7 |
| 0.743872 | 6249.7 |
| 0.751871 | 6249.69 |
| 0.759869 | 6249.69 |
| 0.767868 | 6249.68 |
| 0.775867 | 6249.68 |
| 0.783865 | 6249.68 |
| 0.791864 | 6249.67 |
| 0.799862 | 6249.67 |
| 0.807861 | 6249.67 |
| 0.81586 | 6249.66 |
| 0.823858 | 6249.66 |
| 0.831857 | 6249.65 |
| 0.839855 | 6249.65 |
| 0.847854 | 6249.65 |
| 0.855853 | 6249.64 |
| 0.863851 | 6249.64 |
| 0.87185 | 6249.63 |
| 0.879848 | 6249.63 |
| 0.887847 | 6249.62 |
| 0.895846 | 6249.62 |
| 0.903844 | 6249.62 |
| 0.911843 | 6249.61 |
| 0.919841 | 6249.61 |
| 0.92784 | 6249.6 |
| 0.935839 | 6249.6 |
| 0.943837 | 6249.59 |
| 0.951836 | 6249.59 |
| 0.959834 | 6249.58 |
| 0.967833 | 6249.58 |
| 0.975832 | 6249.57 |
| 0.98383 | 6249.57 |
| 0.991829 | 6249.56 |
| 0.999827 | 6249.56 |
| 1.00783 | 6249.56 |
| 1.01582 | 6249.55 |
| 1.02382 | 6249.54 |
| 1.03182 | 6249.54 |
| 1.03982 | 6249.53 |
| 1.04782 | 6249.53 |
| 1.05582 | 6249.52 |
| 1.06382 | 6249.52 |
| 1.07182 | 6249.51 |
| 1.07981 | 6249.51 |
| 1.08781 | 6249.5 |
| 1.09581 | 6249.5 |
| 1.10381 | 6249.49 |
| 1.11181 | 6249.48 |
| 1.11981 | 6249.48 |
| 1.12781 | 6249.47 |
| 1.1358 | 6249.47 |
| 1.1438 | 6249.46 |
| 1.1518 | 6249.45 |
| 1.1598 | 6249.45 |
| 1.1678 | 6249.44 |
| 1.1758 | 6249.44 |
| 1.1838 | 6249.43 |
| 1.19179 | 6249.42 |
| 1.19979 | 6249.42 |
| 1.20779 | 6249.41 |
| 1.21579 | 6249.4 |
| 1.22379 | 6249.4 |
| 1.23179 | 6249.39 |
| 1.23979 | 6249.38 |
| 1.24778 | 6249.38 |
| 1.25578 | 6249.37 |
| 1.26378 | 6249.36 |
| 1.27178 | 6249.36 |
| 1.27978 | 6249.35 |
| 1.28778 | 6249.34 |
| 1.29578 | 6249.34 |
| 1.30377 | 6249.33 |
| 1.31177 | 6249.32 |
| 1.31977 | 6249.32 |
| 1.32777 | 6249.31 |
| 1.33577 | 6249.3 |
| 1.34377 | 6249.29 |
| 1.35177 | 6249.29 |
| 1.35976 | 6249.28 |
| 1.36776 | 6249.27 |
| 1.37576 | 6249.26 |
| 1.38376 | 6249.26 |
| 1.39176 | 6249.25 |
| 1.39976 | 6249.24 |
| 1.40776 | 6249.23 |
| 1.41576 | 6249.22 |
| 1.42375 | 6249.22 |
| 1.43175 | 6249.21 |
| 1.43975 | 6249.2 |
| 1.44775 | 6249.19 |
| 1.45575 | 6249.18 |
| 1.46375 | 6249.18 |
| 1.47175 | 6249.17 |
| 1.47974 | 6249.16 |
| 1.48774 | 6249.15 |
| 1.49574 | 6249.14 |
| 1.50374 | 6249.13 |
| 1.51174 | 6249.12 |
| 1.51974 | 6249.12 |
| 1.52774 | 6249.11 |
| 1.53574 | 6249.1 |
| 1.54373 | 6249.09 |
| 1.55173 | 6249.08 |
| 1.55973 | 6249.07 |
| 1.56773 | 6249.06 |
| 1.57573 | 6249.05 |
| 1.58373 | 6249.04 |
| 1.59173 | 6249.03 |

| **Visual illusions** | | |
| --- | --- | --- |
| **Black Hole Approach (BHA)** | | |
| 0s | A dark night, Aircraft flying horizontally and leveled, Aircraft at 2000 ft (609.6 m) | Latitude difference = 0.0866 (9616 m), Longitude difference = 0.0434 (4496 m), Aircraft's starting heading = 240°, Landing strip heading = 175° |
| 4s | ATC message: land on landing strip 18 |  |
| SD window – the entire flight duration,  SD response: Y position > 4000m and Altitude < 60 m or a crash Indicates a shallow approach to landing strip | | |
| Visual timeline | | Aircraft and landing strip location |
|  | | Longitude: 0.0434  Longitude: 0  Latitude: 0  Latitude: 0.0866  240  175 |

| Time [s] | Altitude [m] |
| --- | --- |
| 0 | 609.6 |
| 0.00799894 | 609.6 |
| 0.0159976 | 611.051 |
| 0.0239964 | 611.051 |
| 0.0319951 | 611.051 |
| 0.039994 | 611.05 |
| 0.0479925 | 611.05 |
| 0.0559912 | 611.05 |
| 0.0639896 | 611.049 |
| 0.0719883 | 611.049 |
| 0.0799868 | 611.048 |
| 0.0879855 | 611.048 |
| 0.095984 | 611.047 |
| 0.103983 | 611.334 |
| 0.111981 | 611.334 |
| 0.11998 | 611.333 |
| 0.127978 | 611.332 |
| 0.135977 | 611.331 |
| 0.143975 | 611.329 |
| 0.151974 | 611.328 |
| 0.159973 | 611.327 |
| 0.167971 | 611.325 |
| 0.17597 | 611.324 |
| 0.183968 | 611.322 |
| 0.191967 | 612.044 |
| 0.199965 | 612.043 |
| 0.207964 | 612.041 |
| 0.215963 | 612.039 |
| 0.223961 | 612.037 |
| 0.23196 | 612.511 |
| 0.239958 | 612.509 |
| 0.247957 | 612.507 |
| 0.255955 | 612.505 |
| 0.263954 | 612.503 |
| 0.271953 | 612.5 |
| 0.279951 | 613.017 |
| 0.28795 | 613.015 |
| 0.295949 | 613.012 |
| 0.303947 | 613.01 |
| 0.311946 | 613.007 |
| 0.319944 | 613.004 |
| 0.327943 | 613.594 |
| 0.335941 | 613.591 |
| 0.34394 | 613.588 |
| 0.351938 | 613.585 |
| 0.359937 | 613.582 |
| 0.367936 | 613.578 |
| 0.375934 | 614.46 |
| 0.383933 | 614.457 |
| 0.391931 | 614.453 |
| 0.39993 | 614.45 |
| 0.407928 | 614.446 |
| 0.415927 | 614.443 |
| 0.423926 | 614.439 |
| 0.431924 | 614.98 |
| 0.439923 | 614.977 |
| 0.447921 | 614.973 |
| 0.45592 | 614.969 |
| 0.463918 | 614.965 |
| 0.471917 | 614.961 |
| 0.479916 | 615.702 |
| 0.487914 | 615.698 |
| 0.495913 | 615.693 |
| 0.503911 | 615.689 |
| 0.51191 | 615.685 |
| 0.519909 | 615.68 |
| 0.527907 | 616.08 |
| 0.535906 | 616.075 |
| 0.543904 | 616.07 |
| 0.551903 | 616.065 |
| 0.559901 | 616.06 |
| 0.5679 | 616.055 |
| 0.575899 | 616.436 |
| 0.583897 | 616.431 |
| 0.591896 | 616.426 |
| 0.599894 | 616.42 |
| 0.607893 | 616.415 |
| 0.615891 | 616.409 |
| 0.62389 | 616.404 |
| 0.631889 | 616.849 |
| 0.639887 | 616.843 |
| 0.647886 | 616.837 |
| 0.655884 | 616.831 |
| 0.663883 | 616.825 |
| 0.671881 | 616.819 |
| 0.67988 | 617.274 |
| 0.687879 | 617.268 |
| 0.695877 | 617.261 |
| 0.703876 | 617.255 |
| 0.711874 | 617.248 |
| 0.719873 | 617.242 |
| 0.727871 | 617.978 |
| 0.73587 | 617.971 |
| 0.743869 | 617.965 |
| 0.751867 | 617.958 |
| 0.759866 | 617.951 |
| 0.767864 | 617.944 |
| 0.775863 | 618.68 |
| 0.783862 | 618.672 |
| 0.79186 | 618.665 |
| 0.799859 | 618.658 |
| 0.807857 | 618.65 |
| 0.815856 | 618.642 |
| 0.823854 | 618.635 |
| 0.831853 | 619.376 |
| 0.839852 | 619.368 |
| 0.84785 | 619.36 |
| 0.855849 | 619.352 |
| 0.863847 | 619.344 |
| 0.871846 | 619.336 |
| 0.879844 | 620.052 |
| 0.887843 | 620.044 |
| 0.895842 | 620.035 |
| 0.90384 | 620.026 |
| 0.911839 | 620.018 |
| 0.919837 | 620.009 |
| 0.927836 | 620.638 |
| 0.935834 | 620.63 |
| 0.943833 | 620.62 |
| 0.951832 | 620.611 |
| 0.95983 | 620.602 |
| 0.967829 | 620.593 |
| 0.975827 | 621.583 |
| 0.983826 | 621.574 |
| 0.991825 | 621.564 |
| 0.999823 | 621.555 |
| 1.00782 | 621.555 |
| 1.01582 | 621.535 |
| 1.02382 | 621.525 |
| 1.03182 | 622.563 |
| 1.03982 | 622.553 |
| 1.04782 | 622.543 |
| 1.05581 | 622.533 |
| 1.06381 | 622.522 |
| 1.07181 | 622.512 |
| 1.07981 | 623.566 |
| 1.08781 | 623.555 |
| 1.09581 | 623.545 |
| 1.10381 | 623.534 |
| 1.1118 | 623.523 |
| 1.1198 | 623.511 |
| 1.1278 | 624.318 |
| 1.1358 | 624.307 |
| 1.1438 | 624.296 |
| 1.1518 | 624.284 |
| 1.1598 | 624.273 |
| 1.16779 | 624.261 |
| 1.17579 | 625.188 |
| 1.18379 | 625.176 |
| 1.19179 | 625.164 |
| 1.19979 | 625.152 |
| 1.20779 | 625.14 |
| 1.21579 | 625.128 |
| 1.22378 | 625.115 |
| 1.23178 | 625.916 |
| 1.23978 | 625.903 |
| 1.24778 | 625.89 |
| 1.25578 | 625.877 |
| 1.26378 | 625.864 |
| 1.27178 | 625.851 |
| 1.27977 | 626.732 |
| 1.28777 | 626.718 |
| 1.29577 | 626.705 |
| 1.30377 | 626.691 |
| 1.31177 | 626.678 |
| 1.31977 | 626.664 |
| 1.32777 | 627.416 |
| 1.33576 | 627.402 |
| 1.34376 | 627.388 |
| 1.35176 | 627.374 |
| 1.35976 | 627.359 |
| 1.36776 | 627.345 |
| 1.37576 | 628.096 |
| 1.38376 | 628.081 |
| 1.39175 | 628.067 |
| 1.39975 | 628.052 |
| 1.40775 | 628.037 |
| 1.41575 | 628.022 |
| 1.42375 | 628.007 |
| 1.43175 | 628.757 |
| 1.43975 | 628.742 |
| 1.44774 | 628.726 |
| 1.45574 | 628.711 |
| 1.46374 | 628.695 |
| 1.47174 | 628.679 |
| 1.47974 | 629.556 |
| 1.48774 | 629.54 |
| 1.49574 | 629.524 |
| 1.50373 | 629.508 |
| 1.51173 | 629.492 |
| 1.51973 | 629.475 |
| 1.52773 | 630.479 |
| 1.53573 | 630.463 |
| 1.54373 | 630.446 |
| 1.55173 | 630.43 |
| 1.55973 | 630.413 |
| 1.56772 | 630.396 |
| 1.57572 | 631.47 |
| 1.58372 | 631.453 |
| 1.59172 | 631.436 |

| **Ground-Light and Stars** | | |
| --- | --- | --- |
| 0s | A dark night, Aircraft flying horizontally and leveled, Aircraft altitude = 8300 ft, heading point = 105° |  |
| 20s | ATC message 1 to turn left towards heading point 300 |  |
| Condition | Heading > 330° | Past 0 heading point to 359 ° |
| +0.01s | ATC message 2: maintain height and direction |  |
| +4s | HUD and instruments panel turn black |  |
| SD window time,  SD response: Altitude±152 m, or Air Speed±26 m/s  Indicates a motion towards original roll | | |
| +20s | HUD and instruments panel return to normal |  |
| Visual timeline | | |

Heading poistion [deg]

| Time [s] | aeromodel.headingPos (deg) |
| --- | --- |
| 0 | 105.224 |
| 0.0079987 | 105.224 |
| 0.0159972 | 105.224 |
| 0.0239959 | 105.224 |
| 0.0319943 | 105.224 |
| 0.0399928 | 105.224 |
| 0.0479915 | 105.224 |
| 0.05599 | 105.224 |
| 0.0639887 | 105.224 |
| 0.0719872 | 105.224 |
| 0.0799859 | 105.224 |
| 0.0879843 | 105.224 |
| 0.095983 | 105.224 |
| 0.103981 | 105.224 |
| 0.11198 | 105.224 |
| 0.119979 | 105.224 |
| 0.127978 | 105.224 |
| 0.135977 | 105.224 |
| 0.143975 | 105.224 |
| 0.151974 | 105.224 |
| 0.159973 | 105.224 |
| 0.167971 | 105.224 |
| 0.17597 | 105.224 |
| 0.183968 | 105.224 |
| 0.191967 | 105.224 |
| 0.199965 | 105.224 |
| 0.207964 | 105.224 |
| 0.215963 | 105.224 |
| 0.223961 | 105.224 |
| 0.23196 | 105.224 |
| 0.239958 | 105.224 |
| 0.247957 | 105.223 |
| 0.255955 | 105.223 |
| 0.263954 | 105.223 |
| 0.271953 | 105.223 |
| 0.279951 | 105.223 |
| 0.28795 | 105.223 |
| 0.295948 | 105.223 |
| 0.303947 | 105.223 |
| 0.311945 | 105.223 |
| 0.319944 | 105.223 |
| 0.327943 | 105.223 |
| 0.335941 | 105.223 |
| 0.34394 | 105.223 |
| 0.351938 | 105.223 |
| 0.359937 | 105.223 |
| 0.367936 | 105.223 |
| 0.375934 | 105.223 |
| 0.383933 | 105.223 |
| 0.391931 | 105.223 |
| 0.39993 | 105.223 |
| 0.407928 | 105.223 |
| 0.415927 | 105.223 |
| 0.423926 | 105.223 |
| 0.431924 | 105.222 |
| 0.439923 | 105.222 |
| 0.447921 | 105.222 |
| 0.45592 | 105.222 |
| 0.463918 | 105.222 |
| 0.471917 | 105.222 |
| 0.479916 | 105.222 |
| 0.487914 | 105.222 |
| 0.495913 | 105.222 |
| 0.503911 | 105.222 |
| 0.51191 | 105.222 |
| 0.519908 | 105.222 |
| 0.527907 | 105.222 |
| 0.535906 | 105.222 |
| 0.543904 | 105.222 |
| 0.551903 | 105.222 |
| 0.559901 | 105.222 |
| 0.5679 | 105.222 |
| 0.575898 | 105.222 |
| 0.583897 | 105.222 |
| 0.591896 | 105.222 |
| 0.599894 | 105.222 |
| 0.607893 | 105.222 |
| 0.615891 | 105.222 |
| 0.62389 | 105.222 |
| 0.631889 | 105.222 |
| 0.639887 | 105.222 |
| 0.647886 | 105.222 |
| 0.655884 | 105.222 |
| 0.663883 | 105.222 |
| 0.671881 | 105.222 |
| 0.67988 | 105.222 |
| 0.687879 | 105.222 |
| 0.695877 | 105.222 |
| 0.703876 | 105.222 |
| 0.711874 | 105.222 |
| 0.719873 | 105.222 |
| 0.727871 | 105.222 |
| 0.73587 | 105.222 |
| 0.743869 | 105.222 |
| 0.751867 | 105.222 |
| 0.759866 | 105.222 |
| 0.767864 | 105.222 |
| 0.775863 | 105.222 |
| 0.783861 | 105.222 |
| 0.79186 | 105.221 |
| 0.799859 | 105.221 |
| 0.807857 | 105.221 |
| 0.815856 | 105.221 |
| 0.823854 | 105.221 |
| 0.831853 | 105.221 |
| 0.839852 | 105.221 |
| 0.84785 | 105.221 |
| 0.855849 | 105.221 |
| 0.863847 | 105.221 |
| 0.871846 | 105.221 |
| 0.879844 | 105.221 |
| 0.887843 | 105.221 |
| 0.895842 | 105.221 |
| 0.90384 | 105.221 |
| 0.911839 | 105.221 |
| 0.919837 | 105.221 |
| 0.927836 | 105.221 |
| 0.935834 | 105.221 |
| 0.943833 | 105.221 |
| 0.951832 | 105.221 |
| 0.95983 | 105.221 |
| 0.967829 | 105.221 |
| 0.975827 | 105.221 |
| 0.983826 | 105.221 |
| 0.991824 | 105.221 |
| 0.999823 | 105.221 |
| 1.00782 | 105.221 |
| 1.01582 | 105.221 |
| 1.02382 | 105.221 |
| 1.03182 | 105.221 |
| 1.03982 | 105.221 |
| 1.04781 | 105.221 |
| 1.05581 | 105.221 |
| 1.06381 | 105.221 |
| 1.07181 | 105.221 |
| 1.07981 | 105.221 |
| 1.08781 | 105.221 |
| 1.09581 | 105.221 |
| 1.1038 | 105.221 |
| 1.1118 | 105.221 |
| 1.1198 | 105.221 |
| 1.1278 | 105.221 |
| 1.1358 | 105.221 |
| 1.1438 | 105.221 |
| 1.1518 | 105.221 |
| 1.1598 | 105.221 |
| 1.16779 | 105.221 |
| 1.17579 | 105.221 |
| 1.18379 | 105.221 |
| 1.19179 | 105.221 |
| 1.19979 | 105.221 |
| 1.20779 | 105.221 |
| 1.21579 | 105.221 |
| 1.22378 | 105.221 |
| 1.23178 | 105.221 |
| 1.23978 | 105.221 |
| 1.24778 | 105.221 |
| 1.25578 | 105.221 |
| 1.26378 | 105.221 |
| 1.27178 | 105.221 |
| 1.27977 | 105.221 |
| 1.28777 | 105.221 |
| 1.29577 | 105.221 |
| 1.30377 | 105.221 |
| 1.31177 | 105.221 |
| 1.31977 | 105.221 |
| 1.32777 | 105.221 |
| 1.33576 | 105.221 |
| 1.34376 | 105.221 |
| 1.35176 | 105.221 |
| 1.35976 | 105.221 |
| 1.36776 | 105.221 |
| 1.37576 | 105.221 |
| 1.38376 | 105.221 |
| 1.39175 | 105.221 |
| 1.39975 | 105.221 |
| 1.40775 | 105.221 |
| 1.41575 | 105.221 |
| 1.42375 | 105.221 |
| 1.43175 | 105.221 |
| 1.43975 | 105.221 |
| 1.44774 | 105.221 |
| 1.45574 | 105.221 |
| 1.46374 | 105.221 |
| 1.47174 | 105.221 |
| 1.47974 | 105.221 |
| 1.48774 | 105.221 |
| 1.49574 | 105.221 |
| 1.50373 | 105.221 |
| 1.51173 | 105.221 |
| 1.51973 | 105.221 |
| 1.52773 | 105.221 |
| 1.53573 | 105.221 |
| 1.54373 | 105.221 |
| 1.55173 | 105.221 |
| 1.55972 | 105.221 |
| 1.56772 | 105.221 |
| 1.57572 | 105.221 |
| 1.58372 | 105.221 |
| 1.59172 | 105.221 |

| **False Horizon (FH) – only for aircrew members** | | |
| --- | --- | --- |
| 0s | Aircraft flying horizontally and leveled, Aircraft heading towards 285°, altitude = 2,000 ft (609.6 m) |  |
| 10s | ATC message 1: climb to altitude 4000 ft (1219.2 m) |  |
| 24s | ATC message 2: turn towards heading point 120 | Turn while climbing |
| Condition 1 | Altitude > 3800 ft (1158.2 m) |  |
| +4s | ATC message 3: climb to altitude 9000 ft (2743 m) |  |
| Condition 2 | Altitude > 4480 ft (1365.5 m) |  |
| +0.01s | Cirrus clouds appear, 4/8 coverage, slope 4° | Reduces visual cues |
| Condition 3 | Altitude > 8300 ft (2529.8 m) | Before leaving the clouds HUD artificial horizon disappears |
| +0.01s | Artificial horizon disappears |  |
| Condition 4 | Altitude > 8500 ft (2590.8 m) |  |
| +0.01s | Cirrus clouds appear (4/8 coverage, slope 8°) | Cloud line is at 8° slope |
| +0.8s | ATC message 4: straighten aircraft |  |
| SD window time,  SD response: Roll position <= -5° & Roll Velocity <=0°/s  Indicates a straightening according to the cloud's slope | | |
| +16.5s | HUD returns to normal |  |
| Visual timeline | | |

Altitude [m]

| Time [s] | Altitude [m] |
| --- | --- |
| 0 | 609.6 |
| 0.00799894 | 609.6 |
| 0.0159976 | 611.051 |
| 0.0239964 | 611.051 |
| 0.0319951 | 611.051 |
| 0.039994 | 611.051 |
| 0.0479927 | 611.051 |
| 0.0559914 | 611.052 |
| 0.0639904 | 611.052 |
| 0.0719891 | 611.052 |
| 0.0799878 | 611.052 |
| 0.0879867 | 611.053 |
| 0.0959854 | 611.053 |
| 0.103984 | 611.054 |
| 0.111983 | 611.054 |
| 0.119982 | 611.055 |
| 0.12798 | 611.055 |
| 0.135979 | 611.056 |
| 0.143978 | 611.056 |
| 0.151977 | 611.057 |
| 0.159976 | 611.057 |
| 0.167974 | 611.058 |
| 0.175973 | 611.059 |
| 0.183972 | 611.06 |
| 0.191971 | 611.06 |
| 0.19997 | 611.061 |
| 0.207968 | 611.062 |
| 0.215967 | 611.063 |
| 0.223966 | 611.064 |
| 0.231965 | 611.065 |
| 0.239963 | 611.066 |
| 0.247962 | 611.067 |
| 0.255961 | 611.068 |
| 0.26396 | 611.069 |
| 0.271958 | 611.07 |
| 0.279957 | 611.071 |
| 0.287956 | 611.072 |
| 0.295955 | 611.073 |
| 0.303954 | 611.075 |
| 0.311952 | 611.076 |
| 0.319951 | 611.077 |
| 0.32795 | 611.079 |
| 0.335949 | 611.08 |
| 0.343947 | 611.081 |
| 0.351946 | 611.083 |
| 0.359945 | 611.084 |
| 0.367944 | 611.085 |
| 0.375942 | 611.087 |
| 0.383941 | 611.088 |
| 0.39194 | 611.09 |
| 0.399939 | 611.092 |
| 0.407938 | 611.093 |
| 0.415936 | 611.095 |
| 0.423935 | 611.096 |
| 0.431934 | 611.098 |
| 0.439933 | 611.1 |
| 0.447932 | 611.101 |
| 0.45593 | 611.103 |
| 0.463929 | 611.105 |
| 0.471928 | 611.107 |
| 0.479927 | 611.109 |
| 0.487925 | 611.11 |
| 0.495924 | 611.112 |
| 0.503923 | 611.114 |
| 0.511922 | 611.116 |
| 0.519921 | 611.118 |
| 0.527919 | 611.12 |
| 0.535918 | 611.122 |
| 0.543917 | 611.124 |
| 0.551916 | 611.126 |
| 0.559914 | 611.128 |
| 0.567913 | 611.13 |
| 0.575912 | 611.132 |
| 0.583911 | 611.134 |
| 0.591909 | 611.137 |
| 0.599908 | 611.139 |
| 0.607907 | 611.141 |
| 0.615906 | 611.143 |
| 0.623905 | 611.145 |
| 0.631903 | 611.148 |
| 0.639902 | 611.15 |
| 0.647901 | 611.152 |
| 0.6559 | 611.155 |
| 0.663898 | 611.157 |
| 0.671897 | 611.159 |
| 0.679896 | 611.162 |
| 0.687895 | 611.164 |
| 0.695894 | 611.167 |
| 0.703892 | 611.169 |
| 0.711891 | 611.172 |
| 0.71989 | 611.174 |
| 0.727889 | 611.177 |
| 0.735888 | 611.179 |
| 0.743886 | 611.182 |
| 0.751885 | 611.185 |
| 0.759884 | 611.187 |
| 0.767883 | 611.19 |
| 0.775881 | 611.192 |
| 0.78388 | 611.195 |
| 0.791879 | 611.198 |
| 0.799878 | 611.201 |
| 0.807876 | 611.203 |
| 0.815875 | 611.206 |
| 0.823874 | 611.209 |
| 0.831873 | 611.212 |
| 0.839872 | 611.214 |
| 0.84787 | 611.217 |
| 0.855869 | 611.22 |
| 0.863868 | 611.223 |
| 0.871867 | 611.226 |
| 0.879865 | 611.229 |
| 0.887864 | 611.232 |
| 0.895863 | 611.235 |
| 0.903862 | 611.238 |
| 0.91186 | 611.241 |
| 0.919859 | 611.244 |
| 0.927858 | 611.247 |
| 0.935857 | 611.25 |
| 0.943856 | 611.253 |
| 0.951854 | 611.256 |
| 0.959853 | 611.259 |
| 0.967852 | 611.262 |
| 0.975851 | 611.265 |
| 0.98385 | 611.269 |
| 0.991848 | 611.272 |
| 0.999847 | 611.275 |
| 1.00785 | 611.278 |
| 1.01584 | 611.281 |
| 1.02384 | 611.285 |
| 1.03184 | 611.288 |
| 1.03984 | 611.291 |
| 1.04784 | 611.294 |
| 1.05584 | 611.298 |
| 1.06384 | 611.301 |
| 1.07184 | 611.305 |
| 1.07983 | 611.308 |
| 1.08783 | 611.311 |
| 1.09583 | 611.315 |
| 1.10383 | 611.318 |
| 1.11183 | 611.322 |
| 1.11983 | 611.325 |
| 1.12783 | 611.329 |
| 1.13583 | 611.332 |
| 1.14383 | 611.336 |
| 1.15182 | 611.339 |
| 1.15982 | 611.343 |
| 1.16782 | 611.346 |
| 1.17582 | 611.35 |
| 1.18382 | 611.353 |
| 1.19182 | 611.357 |
| 1.19982 | 611.361 |
| 1.20782 | 611.364 |
| 1.21581 | 611.368 |
| 1.22381 | 611.372 |
| 1.23181 | 611.375 |
| 1.23981 | 611.379 |
| 1.24781 | 611.383 |
| 1.25581 | 611.387 |
| 1.26381 | 611.39 |
| 1.27181 | 611.394 |
| 1.2798 | 611.398 |
| 1.2878 | 611.402 |
| 1.2958 | 611.406 |
| 1.3038 | 611.409 |
| 1.3118 | 611.413 |
| 1.3198 | 611.417 |
| 1.3278 | 611.421 |
| 1.3358 | 611.425 |
| 1.34379 | 611.429 |
| 1.35179 | 611.433 |
| 1.35979 | 611.437 |
| 1.36779 | 611.441 |
| 1.37579 | 611.445 |
| 1.38379 | 611.449 |
| 1.39179 | 611.453 |
| 1.39979 | 611.457 |
| 1.40778 | 611.461 |
| 1.41578 | 611.465 |
| 1.42378 | 611.469 |
| 1.43178 | 611.473 |
| 1.43978 | 611.477 |
| 1.44778 | 611.481 |
| 1.45578 | 611.485 |
| 1.46378 | 611.49 |
| 1.47177 | 611.494 |
| 1.47977 | 611.498 |
| 1.48777 | 611.502 |
| 1.49577 | 611.506 |
| 1.50377 | 611.511 |
| 1.51177 | 611.515 |
| 1.51977 | 611.519 |
| 1.52777 | 611.523 |
| 1.53576 | 611.528 |
| 1.54376 | 611.532 |
| 1.55176 | 611.536 |
| 1.55976 | 611.541 |
| 1.56776 | 611.545 |
| 1.57576 | 611.549 |
| 1.58376 | 611.554 |
| 1.59176 | 611.558 |
